# Global biogeography of the smallest plankton across ocean depths

**DOI:** 10.1101/2023.01.13.523743

**Authors:** Pedro C. Junger, Hugo Sarmento, Caterina. R. Giner, Mireia Mestre, Marta Sebastián, Xosé Anxelu G. Morán, Javier Arístegui, Susana Agustí, Carlos M. Duarte, Silvia G. Acinas, Ramon Massana, Josep M. Gasol, Ramiro Logares

## Abstract

Tiny ocean plankton (picoplankton) are fundamental for the functioning of the biosphere, but the ecological mechanisms shaping their biogeography are partially understood. Comprehending whether these microorganisms are structured by niche vs. neutral processes is highly relevant in the context of global change. The ecological drivers structuring picoplankton communities differ between prokaryotes and minute eukaryotes (picoeukaryotes) in the global surface ocean: while prokaryotic communities are shaped by a balanced combination of *dispersal, selection*, and *drift*, picoeukaryotic communities are mainly shaped by *dispersal limitation*. Yet, whether or not the relative importance of these processes in structuring picoplankton varies as we dive into the deep ocean was unknown. Here we investigate the mechanisms structuring picoplanktonic communities inhabiting different ocean depths. We analyzed 451 samples from the tropical and subtropical global ocean and the Mediterranean Sea covering the epi- (0-200m), meso- (200- 1,000m), and bathypelagic (1,000-4,000m) depth zones. We found that selection decreased with depth possibly due to lower habitat heterogeneity. In turn, dispersal limitation increased with depth, possibly due to dispersal barriers such as water masses and bottom topography. Picoplankton β-diversity positively correlated with environmental heterogeneity and water mass variability in both the open-ocean and the Mediterranean Sea. However, this relationship tended to be weaker for picoeukaryotes than for prokaryotes. Community patterns were generally more pronounced in the Mediterranean Sea, probably because of its substantial cross-basin environmental heterogeneity and deep-water isolation. Altogether, we found that different combinations of ecological mechanisms shape the biogeography of the smallest members of the ocean microbiome across ocean depths.

## Introduction

The smallest eukaryotes and prokaryotes (picoplankton, 0.2 - 3 µm) play essential roles in the global ocean: from trophic interactions (1) to biogeochemical cycles (2, 3). They account for 57% (∼3.8 Gt C) of the ocean’s biomass (4) and are the main contributors to the taxonomic and functional diversity of the ocean (5–8). Therefore, understanding the mechanisms determining their global biogeography is fundamental to predict how they will respond to environmental changes. Picoplankton abundance, diversity, and composition are relatively well described across ocean depths (9, 10): prokaryotes’ diversity increases with depth (8, 11), while picoeukaryotes’ diversity sharply decreases (12). These depth-related patterns are strongly shaped by gradients in sunlight, temperature, oxygen, and nutrients (8, 11) as well as by physical barriers such as water masses, currents, and fronts (13–16). However, the ecological processes underpinning picoplankton biogeography are only partially understood (17, 18), specially considering different ocean depth zones and geographic scales. Given that the deep ocean is the largest ecosystem on our planet and harbors a massive microbial genetic diversity (19) – responsible for essential global ecosystem services – understanding how these processes shape the microbiota in the understudied and vast deep ocean is particularly important.

The biogeography of organisms are the result of four high-level ecological processes that act in different proportions: selection, dispersal, ecological drift, and diversification (20). *Selection* is a deterministic force emerging from combinations of biotic and abiotic variables that lead to differences in the fitness of individuals of a species and, as a consequence, to changes in community structure. *Selection* can either restrict (homogeneous selection) or promote (heterogeneous selection) the divergence of communities (21). *Dispersal* is the movement of organisms across space and their establishment in new locations, affecting local community assembly by adding individuals from the regional species pool. Dispersal is considered a stochastic process for small plankton as they passively drift with currents (21). Microbial dispersal rates may be high (homogenizing dispersal), moderate, or low (dispersal limitation) (21), depending on organism and population sizes, geographic scale, and the presence of physical barriers (18, 22, 23). *Dispersal limitation* takes place when species are not present in suitable habitats because colonizers cannot reach them (24). Thus, the relative importance of *dispersal limitation* usually increases with geographic scales (25) or barriers (22). Ecological drift (hereafter *drift*) refers to random changes in community structure due to stochastic demographic events (i.e., birth, death, immigration, and emigration) in a local community (20). *Drift* is a stochastic process that tends to be most important for the local extinction of low-abundant microbial taxa with small populations (26), especially under a low dispersal scenario (23). Finally, diversification (also referred to as ‘speciation’) is the emergence of new species by evolution (20), which occurs more frequently for microbes than for larger organisms due to their short generation times, high mutation rates as well as horizontal gene transfer (21, 26). Yet, diversification is expected to have a relatively small impact on the turnover of communities that are highly connected via dispersal (27), as is the case for ocean picoplankton (22). *Diversification*, as measured by the evolution of the rRNA gene sequence, will not be further considered here, given that its impact on measured ecological processes is likely minor considering the low evolutionary rates of this marker (28).

A recent study – using *Malaspina* and TARA data – found that the relative importance of these processes differs between the components of the surface ocean picoplankton community: while prokaryotes are shaped by a balanced combination of dispersal, selection, and drift, picoeukaryotes are mainly driven by dispersal limitation (17). However, we do not fully understand whether these processes change across ocean depth zones. These zones display striking differences in environmental and geographic features that may influence selection, dispersal, and drift. First, environmental heterogeneity – potentially exerting heterogeneous selection on microbial communities (17, 29) – is higher in the upper ocean due to stronger horizontal environmental gradients (30) than in the deep ocean (31). Second, the presence of aerial dispersal (32) and faster oceanic currents likely increases dispersal at the surface (33, 34), while the presence of sharper geographical barriers (e.g. water masses and bottom topography) may limit microbial dispersal in the low-turbulent deep ocean (18, 35, 36). Third, smaller population sizes in the deep ocean (9) may lead to reduced dispersal and increased drift (23), as compared to the surface ocean (17, 34). Recently, using a subset of the *Malaspina* dataset, it has been shown that picoplankton community assembly differed between a water layer in the surface ocean (3 m) and a counterpart in the deep ocean (∼4,000 m), with dispersal limitation being relatively more important in the deep layer than in the surface counterpart (18).

In addition, we do not know whether these processes would be different in an ocean basin presenting strong environmental gradients and obvious geographic barriers. In this regard, the Mediterranean Sea – the largest semi-enclosed sea on Earth – is an ideal ocean model to test ecological hypotheses at a smaller scale (37, 38). Although the Mediterranean Sea is connected to the adjacent Atlantic Ocean through the Strait of Gibraltar, it is so in a rather restricted way (39). As a consequence, the Mediterranean Sea has developed unique oceanographic features in comparison to the open ocean, such as higher temperature and salinity in deep waters as well as a west-to-east gradient of decreasing nutrient concentration and increasing salinity in surface waters (11, 40). Additionally, the Mediterranean Sea deep (> 1,000 m) waters are physically divided by the Sicily Strait (500 m deep) into Western and Eastern basins. These features are expected to influence the processes shaping picoplankton biogeography and, ultimately, be reflected in its community composition (11).

In the last few years, it has been found that different processes shape prokaryotes and picoeukaryotes in the surface ocean (17). In addition, a recent report points to differences in the picoplankton biogeography between specific waters layers in the surface (3 m) and deep ocean (4,000 m) (18). However, we still lacked a broad examination of the ecological processes driving picoplankton community assembly and biogeography across all depth zones of the global ocean that takes into account environmental heterogeneity, potential dispersal barriers, and geography. Here, we addressed the previous challenge. We determined the relative importance of the ecological processes structuring picoplanktonic communities inhabiting three ocean depth zones at the global and basin scales: epi- (0-200 m), meso- (200-1,000 m) and bathypelagic (1,000-4,000 m). We also aimed at understanding to what extent water masses, deep-sea topography as well as environmental heterogeneity are potentially limiting dispersal or exerting selection on the picoplanktonic communities. To do so, we used 16S and 18S rRNA gene amplicon sequence variants (ASV) from both prokaryotes and picoeukaryotes collected during global and regional expeditions covering the tropical and subtropical global ocean as well as the Mediterranean Sea. Overall, we hypothesize that the role of heterogeneous selection will decrease with depth due to a potential decrease in habitat heterogeneity, while homogeneous selection is expected to be higher in the bathypelagic compared to the meso- and epipelagic. In turn, the relative importance of dispersal limitation is expected to increase with depth, given the decrease in current speed in deep waters, the existence of geographical barriers (e.g. fronts, deep sea topography), and the absence of aerial dispersal. We also hypothesize that these patterns should be more pronounced in the Mediterranean Sea due to its strong environmental gradients and constrained communities exchange in deep waters.

## Results

### Different ecological processes shape picoplankton communities in depth zones of the ocean

We analyzed picoplankton community composition in 451 samples across three ocean depth zones: epi- (0-200 m – including the deep chlorophyll maxima, DCM), meso- (200-1,000 m), and bathypelagic (1,000-4,000 m) using metabarcoding of the 16S and 18S rRNA genes (Fig. 1A and *SI Appendix*, Fig. S1*A*; see *Methods* for details on standard protocols). These zones display contrasting environmental features across the water column (Fig. 1B and *SI Appendix*, Fig. S1*B*), reflected in a depth-structured picoplankton community composition (Fig. 1C). Our data also makes evident an inverted diversity pattern between the two main components of the picoplankton community: while prokaryotic diversity (richness, Shannon index, and phylogenetic diversity) increased with depth, picoeukaryotic diversity decreased towards the deep ocean (Fig. 1D and *SI Appendix*, Fig. S2). While the Mediterranean Sea displayed higher temperature and salinity as well as lower nutrients than the oceanic basins, particularly in the meso- and bathypelagic (Fig. 1B), the diversity patterns were similar in both ocean sets. The environmental features, however, were reflected in differences in picoplankton community composition (Bray-Curtis Dissimilarity) between the Mediterranean Sea and the rest of the oceanic basins (Fig. 1C). The Mediterranean Sea was evaluated separately from the open ocean in downstream analyses to test whether the large scale patterns are reflected at the regional scale of a smaller basin with strong environmental gradients and sharp geographic barriers.

**Figure 1.**
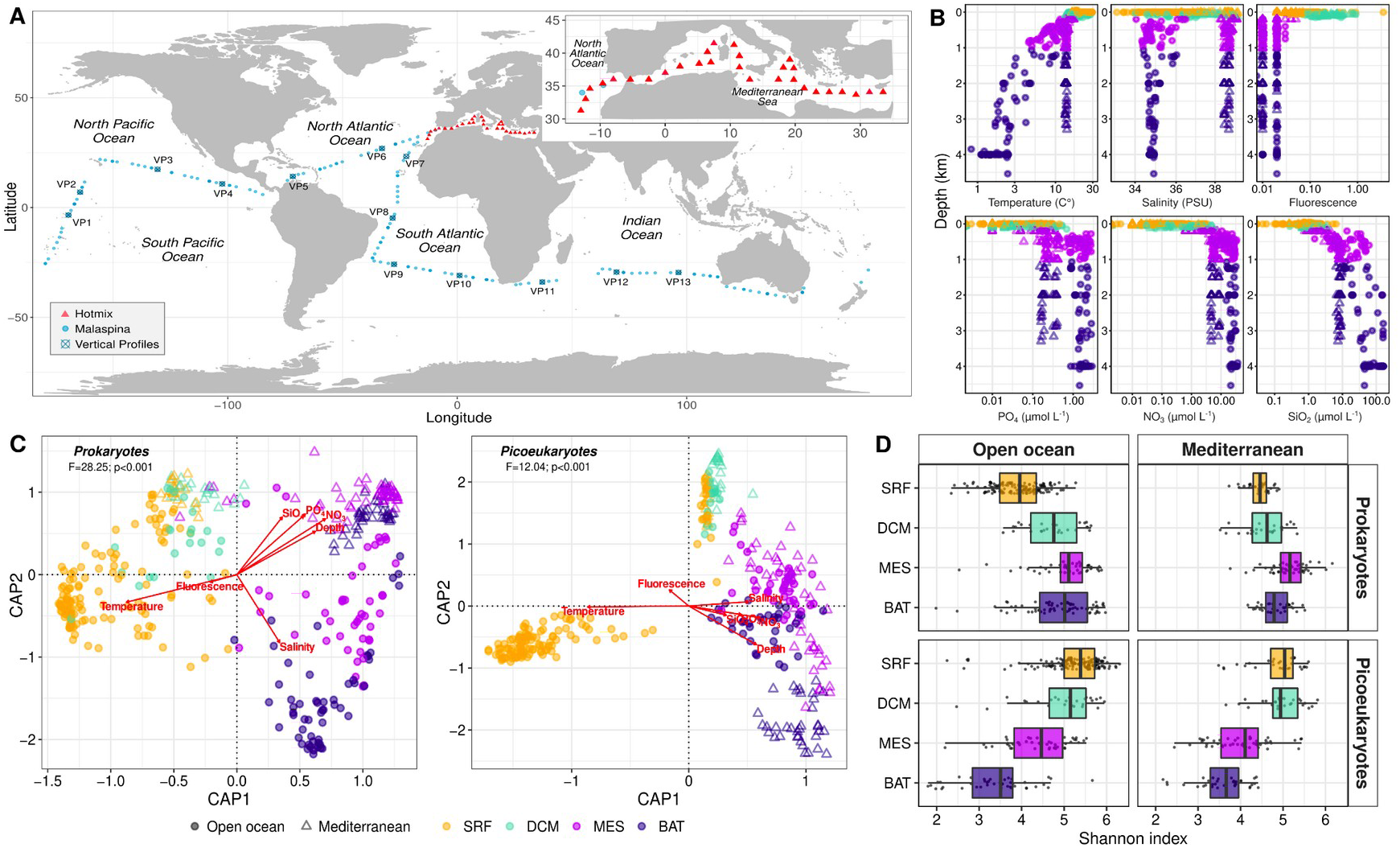
The analyzed dataset covers environmentally and biologically contrasting depth zones of the ocean. **(A)** Geographic distribution of the sampled stations (N=149) from which seawater samples and environmental data were collected at different depth zones (see *SI Appendix*, Fig. S1 for sample vertical distribution) in the two cruises used in this study: *Malaspina-2010* (circumglobal expedition) and *HotMix* (trans-Mediterranean expedition). Stations for which the whole vertical profile was studied in *Malaspina* are represented by crossed squares (13 stations in *Malaspina*). Samples were separated into “open-ocean” (*Malaspina-2010* + *Hotmix* North Atlantic samples) and “Mediterranean Sea” (see reasoning in the *Methods*). **(B)** Vertical profiles of the environmental parameters: temperature, salinity, and fluorescence (Chlorophyll *a* proxy) that decrease with depth, while nutrient concentrations (NO3, PO4, and SiO2) increase with depth. Higher temperature and salinity values and lower nutrient concentrations were observed in the Mediterranean Sea, especially in the meso- and bathypelagic (*SI Appendix*, Fig. S1*B*). **(C)** dbRDA analyses (based on Bray-Curtis dissimilarities) performed on picoplankton community composition of both prokaryotic (left) and picoeukaryotic (right) samples based on 16S rRNA and 18S rRNA genes, respectively. Both communities were structured by depth zones and segregated between the tropical and subtropical open-ocean and the Mediterranean Sea. **(D)** Picoplankton diversity expressed as Shannon index by depth zones (SRF, surface; DCM, deep chlorophyll maxima; MES, Mesopelagic; BAT, Bathypelagic). See Fig. S2 (*SI Appendix)* for picoplankton phylogenetic diversity, gamma diversity, ASVs richness, and Pielou’s evenness index variation by depth zones and correlations with environmental variables.

We found differences in the biodiversity metrics (βNTI, RCBray and β-diversity partitioning *SI Appendix*, Fig. S3 and Fig. S4) and, ultimately, in the balance between ecological processes shaping picoplankton communities across depth zones of the ocean (Fig. 2A). *Selection* explained a similar percentage of the turnover of picoeukaryotes as compared to prokaryotes in the epi- (∼37% vs. ∼36%), meso- (∼32% vs. ∼31%) and bathypelagic (∼32% vs. ∼26%) of the open ocean (Fig. 2A). *Heterogeneous selection* tended to increase with depth for both domains: while for prokaryotes it increased from ∼10% to ∼19% and ∼13% in the meso- and bathypelagic, it increased from ∼13% in the epi-to ∼27% and ∼31% in the meso- and bathypelagic for picoeukaryotes, respectively (Fig. 2A). Accordingly, the relative importance of *homogeneous selection* for prokaryotes decreased from ∼26% in the epi-to ∼13% in the bathypelagic. Similarly, the relative importance of *homogeneous selection* for picoeukaryotes drastically decreased from ∼26 % in the epipelagic to ca. 0.7% in the bathypelagic (Fig. 2A). These patterns were slightly different in the Mediterranean Sea when compared to the tropical and subtropical open ocean. The relative weight of *selection* for the prokaryote community assembly was consistently higher than for the picoeukaryotic counterpart in the epi- (∼54% vs. ∼44%), meso- (∼39% vs. ∼25%) and bathypelagic (∼32 vs. ∼6%, respectively) (Fig. 2A). The proportion of *heterogeneous selection* for prokaryotes dramatically dropped from 37% in the epipelagic to ∼5% in deep waters, while the role of *homogeneous selection* increased from the epi- (∼18%) to the meso- (∼34%) and bathypelagic (∼28%) (Fig. 2A). For picoeukaryotes, both *heterogeneous* and *homogeneous selection* decreased from the epi- (33% and 10%) to the bathypelagic (6% and 0.2%, respectively) (Fig. 2A).

**Figure 2.**
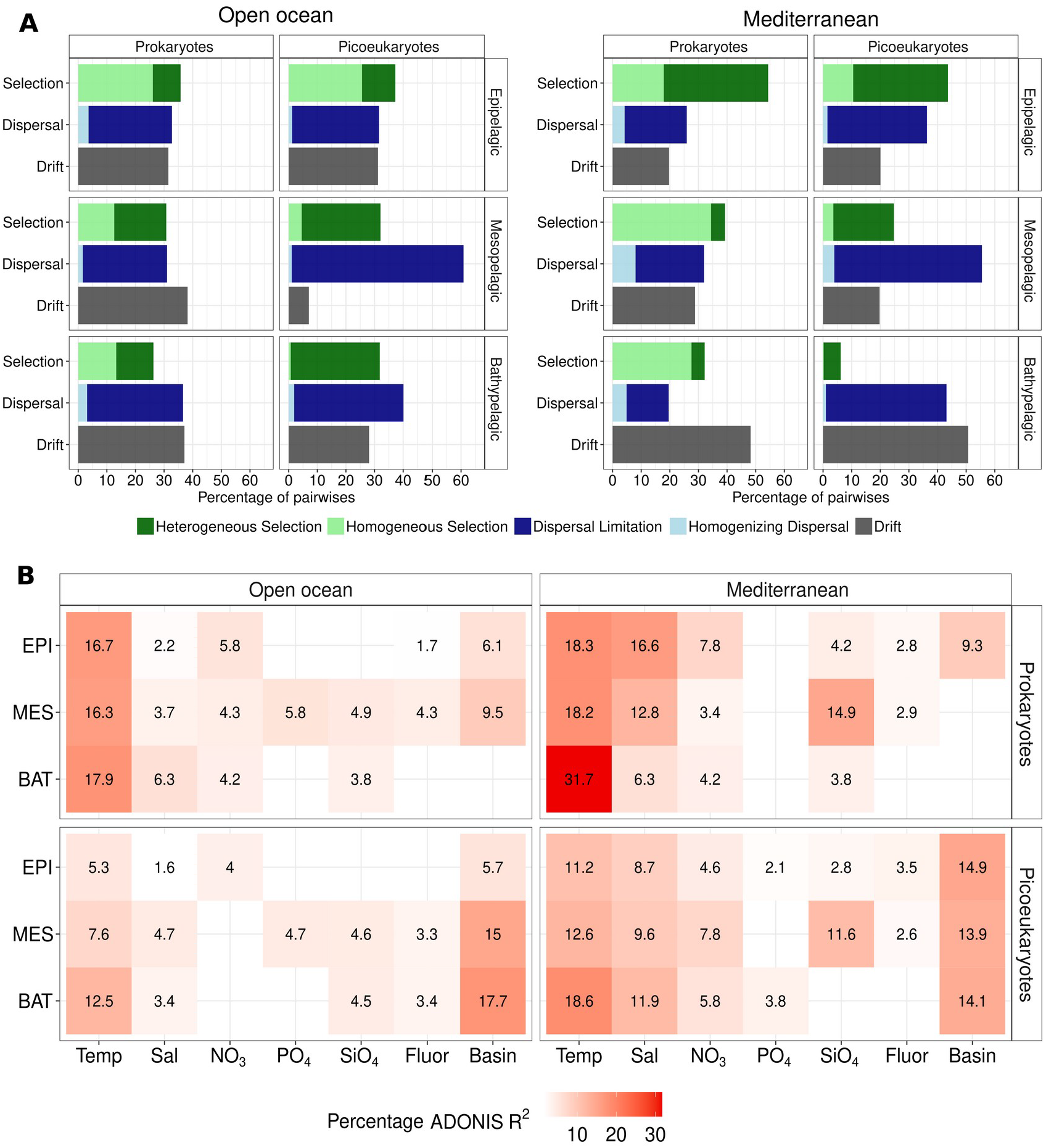
Picoplankton community assembly processes and environmental drivers across ocean depth zones. **(A)** Relative importance of the ecological processes structuring the communities in different depth zones of the global-ocean: Epi- (N=240), Meso- (N=97), and Bathypelagic (N=86). The results with standard evenly-distributed sampling sizes were nearly the same (*SI Appendix*, Fig. S5). The EPI results separated by SRF and DCM are available in the *SI Appendix* (Fig. S7). **(B)** Percentage of variance (Adonis R^2^) in picoeukaryotic and prokaryotic community composition (Bray-Curtis dissimilarity) explained by each environmental variable and ocean basin. Blank spaces depict non-significant results (*p*>0.05). Temp – temperature; Sal – salinity; Fluor – fluorescence; Basin – ocean basin.

### Dispersal limitation

was a more important driver of picoeukaryotic than prokaryotic assembly in the deep zones, especially in the mesopelagic (∼60% vs. ∼29%), of the open ocean. We found that, for picoeukaryotes, the proportion of *dispersal limitation* increased from ∼31 % in the epi-to ∼60% in the meso- and to ∼38% in the bathypelagic (Fig. 2A). In the Mediterranean Sea the relative importance of *dispersal limitation* was much higher for picoeukaryotic than for prokaryotic assembly in the epi- (∼35% vs. ∼22%), meso- (∼52% vs. ∼24%), and bathypelagic (∼42 vs. ∼15%). Conversely, *homogenizing dispersal* had a very limited role in the structuring of the microbiota in all depth zones of the open ocean (<2% for picoeukaryotes and <4% for prokaryotes) and the Mediterranean Sea (<5% for picoeukaryotes and <8% for prokaryotes) (Fig. 2A). *Drift* explained a higher fraction of community turnover for prokaryotes than picoeukaryotes in the meso- (∼38% vs ∼7%) and bathypelagic (∼37% vs ∼28%) of the open ocean (Fig. 2A). This pattern was partially observed in the Mediterranean Sea with *drift* explaining a higher proportion of community turnover for prokaryotes (∼29%) and picoeukaryotes (∼20%) in the mesopelagic (Fig. 2A). While in the open ocean the percentage of turnover explained by *drift* increased with depth for prokaryotes and decreased for picoeukaryotes (Fig. 2A), it sharply increased with depth for both prokaryotes and picoeukaryotes in the Mediterranean Sea (Fig. 2A). When estimated using a standardized sampling-size dataset (N=39 in each depth zone) with evenly-distributed samples (*SI Appendix*, Fig. S5*A* and Fig. S6), the different ecological processes explained fairly similar percentages of variability and the values were strongly linked (R^2^ ∼ 0.9, p<0.001) to those found with the complete dataset (Fig. 2).

When globally estimated (all depths together), *selection* was by far the most relevant ecological process shaping both prokaryotes (∼67%) and picoeukaryotes (∼54%) using both datasets (*SI Appendix*, Fig. S8). *Dispersal limitation* also tended to play a relatively more important role shaping picoeukaryotes than prokaryotes when estimated across all depth zones (*SI Appendix*, Fig. S8). Due to the potential vertical connectivity between the surface and the deep ocean (see detailed reasoning in *Methods*), we also estimated the ecological processes integrating all depths (from 3 to 4,000 m) in each of the 13 vertical profile stations (Fig. 1A). We found that *selection* was consistently the most important factor vertically shaping free-living picoplankton communities in the vertical profile stations, explaining ∼52-81% of the prokaryotic community turnover (*SI Appendix*, Fig. S9) and ∼24-52% of the picoeukaryotic community turnover (*SI Appendix*, Fig. S9). The role of vertical *dispersal limitation* ranged from 10% to 43% in prokaryotes and from 5% to 43% in picoeukaryotes (*SI Appendix*, Fig. S9). The role of *drift* was greater in picoeukaryotes (∼15-43%) than in prokaryotes (∼5-24%) across vertical profiles (*SI Appendix*, Fig. S9).

### The relative importance of selection is ruled by differences in environmental heterogeneity across depth zones

We also evaluated the abiotic drivers of selection across depth zones. Water temperature was the most important environmental driver of prokaryotic community composition in the open ocean (∼16-18%) and the Mediterranean Sea (∼18-32%) (Fig. 2B). Temperature explained a moderate percentage of variance in picoeukaryotic communities inhabiting the bathypelagic (∼12% in the open ocean and ∼18% in the Mediterranean Sea) (Fig. 2B). The percentages of variance in the prokaryotic and picoeukaryotic communities explained by temperature increased from the surface (∼18% and ∼11%, respectively) to the deep zones (∼32% and ∼19%, respectively) in the Mediterranean Sea. Salinity explained a moderate fraction of prokaryotic (up to 16%) and picoeukaryotic (up to 12 %) community variance in the Mediterranean Sea, but not in the open ocean (Fig. 2B). Geography (ocean basin) could affect the structure of picoplankton communities if it is associated with dispersal processes or to different selection regimes exerted in different ocean basins. Geography (ocean basin) explained most of the variation in picoeukaryotes in the open ocean and the Mediterranean Sea (Fig. 2B). Interestingly, the percentage of community variance explained by geography in picoeukaryotes increased markedly from the surface to the meso- and bathypelagic in the open ocean (Fig. 2B). In turn, geography explained a limited fraction of community variance in prokaryotes.

Environmental heterogeneity (average pairwise dissimilarity based on temperature, salinity, fluorescence, PO_4_^3−^, NO_3_^−^, and SiO_2_) was significantly higher in the epi-than in the meso- and bathypelagic of the open ocean and the Mediterranean Sea (*SI Appendix*, Fig. S10). We found that the picoplankton communities’ dissimilarity increased with environmental distance in all depth zones (Fig. 3). This positive relationship was always stronger in the epipelagic than in the bathypelagic (Fig. 3). Prokaryotes displayed a stronger coupling with environmental distance than picoeukaryotes in all depth zones of both the open ocean and the Mediterranean Sea (Fig. 3) and this coupling was stronger in the Mediterranean Sea than in the open ocean across all zones (Fig. 3). When globally estimated (all depth zones together), the community dissimilarity correlation with environmental distance was stronger for prokaryotes than for picoeukaryotes in both the open ocean (r=0.62 *vs*. r=0.46, p<0.001) and the Mediterranean Sea (r=0.69 *vs*. r=0.65, p<0.001) (*SI Appendix*, Fig. S11). The metric used to estimate *selection* (βNTI) was positively correlated, in prokaryotic and picoeukaryotic communities, with environmental distances in both the open ocean (r=0.55 and r=0.50, p<0.001) and the Mediterranean Sea (r=0.55 and r=0.50, p<0.001) (*SI Appendix*, Fig. S11).

**Figure 3.**
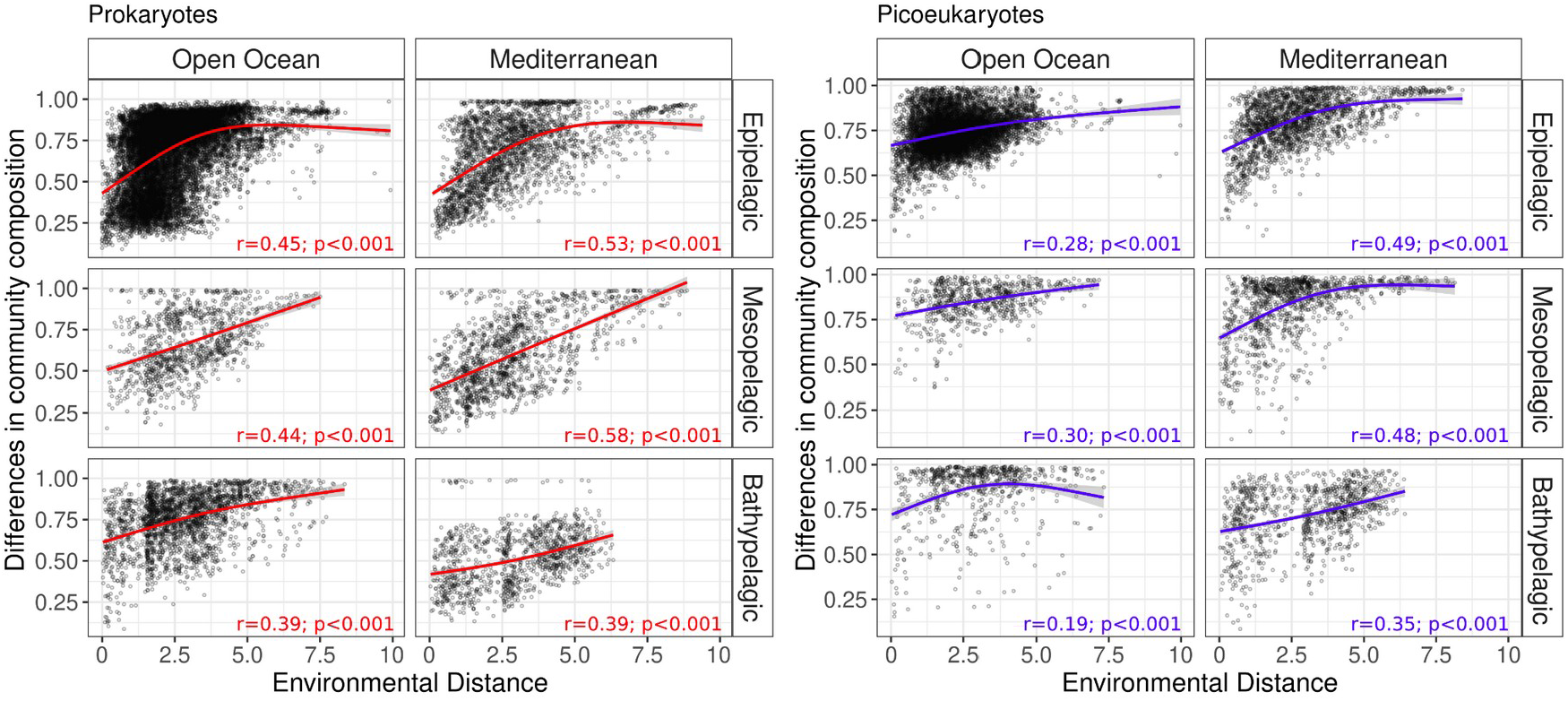
Picoplankton community composition is positively related to environmental heterogeneity. Bray-curtis dissimilarities for all pairwise picoplankton community comparisons as a function of environmental distance for both prokaryotes and picoeukaryotes in the epi-, meso-, and bathypelagic of the open ocean and Mediterranean Sea. The solid curves illustrate the nonlinear regressions. Spearman’s rank correlation coefficients are depicted on the panel. Outliers with high environmental distances (>10) corresponding to pairwise comparisons with epipelagic samples from the Costa Rica Dome upwelling system were removed from the open ocean plot (*SI Appendix*, Fig. S12).

### The role of water masses and deep sea topography in modulating picoplankton assembly

Water masses, which were determined for the meso- and bathypelagic, were vertically structured and segregated by basins in the open ocean and the Mediterranean Sea (*SI Appendix*, Fig. S13). We found that prokaryotic community composition (Bray-Curtis dissimilarity) was positively linked with differences in water mass composition (Euclidean distances) in the meso- and bathypelagic of the open ocean (r=0.2 and r=0.4, p<0.001) and the Mediterranean Sea (r=0.46 and r=0.33, p<0.001) (Fig. 4). For picoeukaryotes, this coupling was generally weaker than for prokaryotes in both the open ocean (r=0.14, p<0.001) and the Mediterranean Sea (r=0.49 and r=0.29, p<0.001) (Fig. 4). In the Mediterranean Sea, the link between picoplankton community and water mass composition was stronger in the meso-than in the bathypelagic (Fig. 4). Strong positive relationships between the picoplankton community and water mass’ composition were also observed within most individual vertical-profile stations, with variable slopes in each station (*SI Appendix*, Fig. S14).

**Figure 4.**
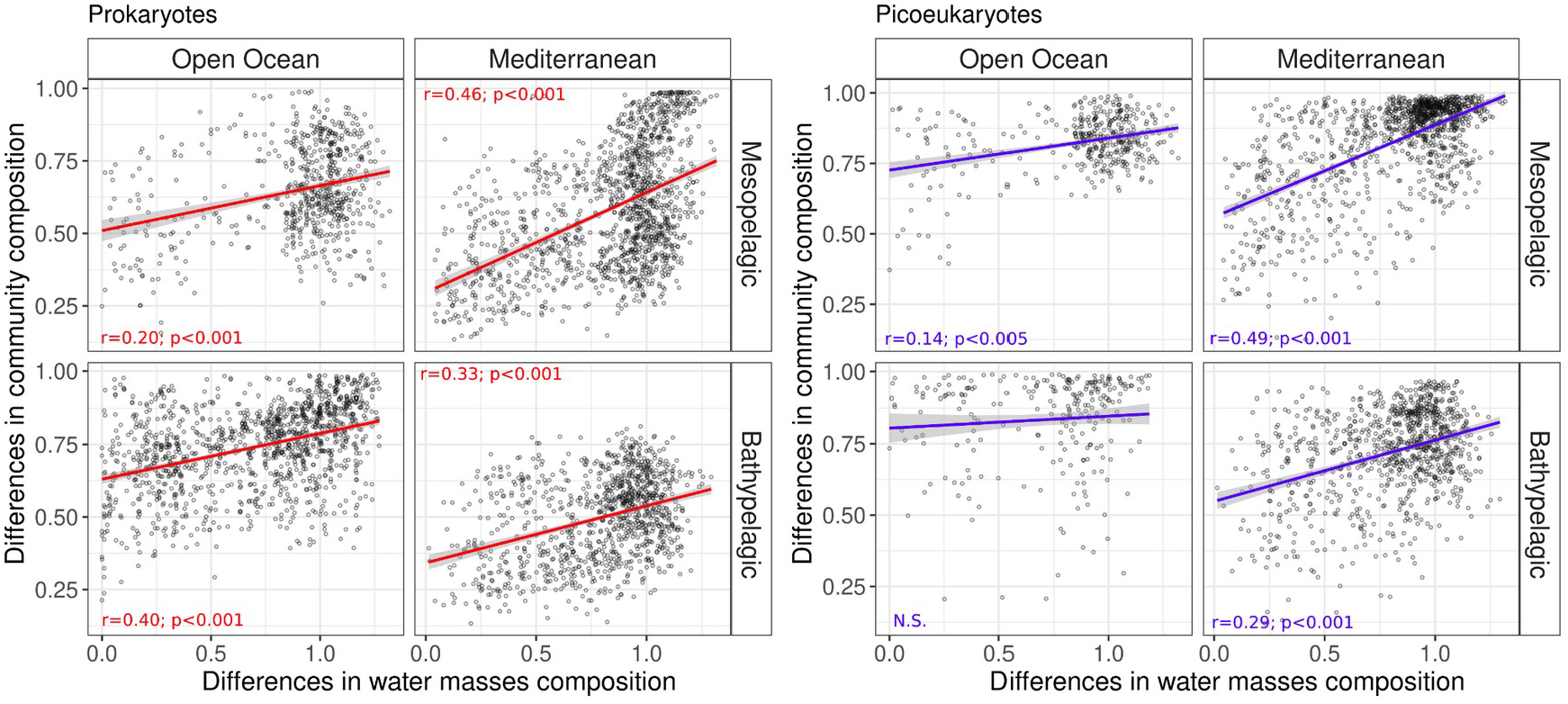
Picoplankton community composition is linked to differences in water mass composition. Bray-Curtis dissimilarity of pairwise picoplankton community comparisons as a function water mass composition dissimilarity (based on euclidean distances) for both prokaryotes and picoeukaryotes in the meso- and bathypelagic of the open ocean and Mediterranean Sea. The solid curves illustrate the nonlinear regressions. Spearman’s rank correlation coefficients are depicted on the panel. N.S.= non significant.

In the open ocean, changes in prokaryotic and picoeukaryotic community composition (β-diversity) displayed positive correlations with geographic distances (distance-decay) in four depth zones (Fig. 5A) even though correlations were weaker for prokaryotes than for picoeukaryotes in most of them. Prokaryotes displayed positive correlations with distances up to ∼2,000 km in the surface and 1,000 km in the deep ocean, while picoeukaryotes showed positive correlations up to ∼3,000 km in the surface and ∼4,000 km in the deep ocean (Fig. 5A). For picoeukaryotes, these positive correlations were stronger in the bathypelagic (Mantel *r* = 0.5, p<0.05) than in the surface (Mantel *r* = 0.3, p<0.05) (Fig. 5A). Interestingly, picoeukaryotes also displayed negative correlations with increasing distances up to ∼ 20,000 km across the deep zones (Fig. 5A). In fact, picoeukaryotes had a higher variation in the spatial autocorrelations than prokaryotes in the deep ocean, especially in the bathypelagic. When evaluating these spatial autocorrelations at a regional scale as in the Mediterranean Sea, we found that prokaryotes and picoeukaryotes did not display such contrasting correlation scores as in the open ocean (Fig. 5A). Indeed, these two domains had similar patterns of positive correlations in the first 350-850 km of the Mediterranean Sea (Fig. 5A). Picoeukaryotes had higher mean sequential changes in communities (β-diversity) than prokaryotes in all depth zones (Fig. 5B). Overall, sequential community change tended to increase with depth in picoeukaryotes, with significant differences between the surface and the meso- and bathypelagic in picoeukaryotes, but not in prokaryotes (Fig. 5B).

**Figure 5.**
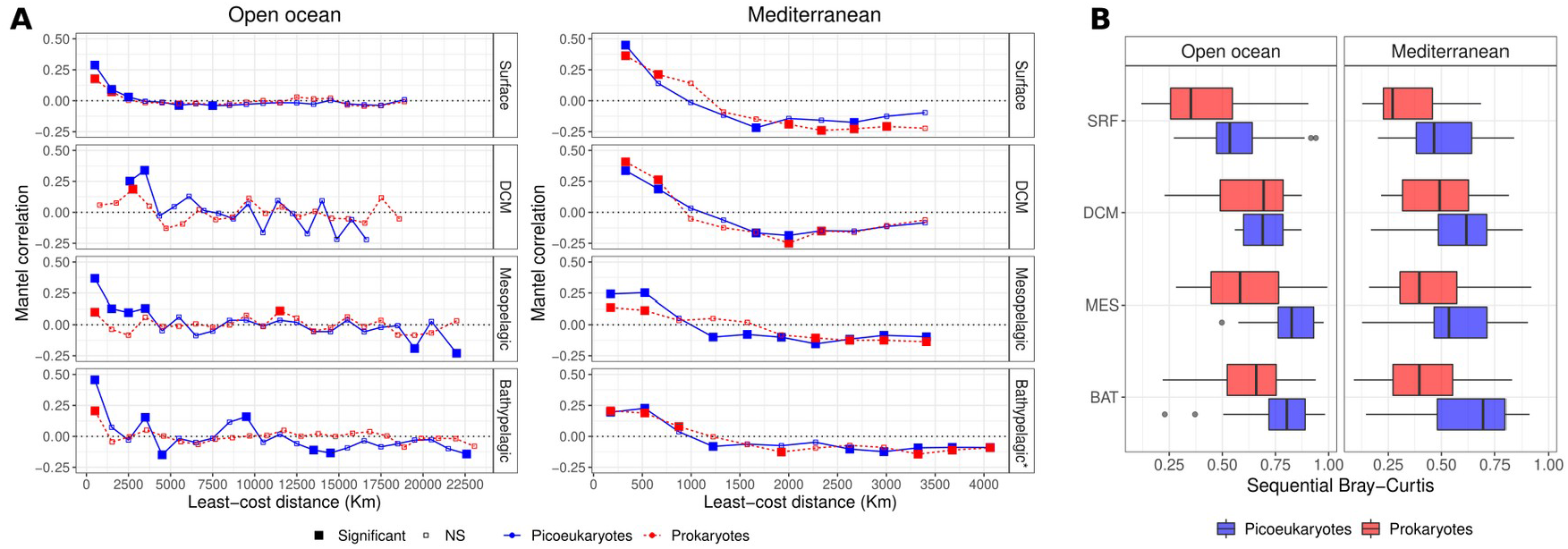
Distance-decay and sequential spatial differentiation in picoplankton communities across ocean depth zones. **(A)** Mantel correlograms between β-diversity and least-cost geographic distances featuring distance classes of 1,000 km for the open ocean and 350 km for the Mediterranean Sea. Filled squares depict significant correlations (*p*<0.05). NS – non-significant correlations. **(B)** Sequential Bray-Curtis dissimilarity values for prokaryotes and picoeukaryotes in all depth zones (means were significantly different between domains [Wilcoxon test, *p*<0.05] in all depth zones, apart from the DCM). The averages were also significantly different (ANOVA, Tukey post-hoc test; *p*<0.001) between the SRF and the deep zones (MES and BAT) for picoeukaryotes, but not for prokaryotes. See Fig. S15 (*SI Appendix*) for maps showing the sequential change in community composition across space in the surface and bathypelagic ocean. The epipelagic was here separated into surface and DCM because we aimed at evaluating only the horizontal geographic distance in each depth.

Microbial abundances and activity may also work as potential regulators of dispersal limitation and drift. Here, microbial abundances – as measured by flow-cytometry – sharply decreased with depth in both the open ocean and the Mediterranean Sea (*SI Appendix*, Fig. S16*A*). Similarly, prokaryotic activity – as measured by leucine incorporation rates – drastically decreased from surface to deep ocean waters (*SI Appendix*, Fig. S16*B*), with statistically significant differences between epipelagic (SRF and DCM) and deep zones (MES and BAT).

## Discussion

### Selection decreases while dispersal limitation and drift increase with depth

Our results support our main hypothesis, indicating that a different combination of ecological processes shapes picoplankton biogeography across ocean depth zones at global and regional scales (Fig. 6). *Selection* was the most important process shaping picoplankton in the epipelagic ocean (see also *SI Appendix, SI Discussion*), likely as a response to a higher overall environmental heterogeneity when compared to the deep ocean. In particular, microalgal blooms (30, 41), magnitude of the DCM (8, 42), ocean fronts and eddies (13, 16, 43, 44), and differences in physicochemical variables (Fig. 1B), altogether increase environmental heterogeneity in the upper ocean (Fig. 6). In the epipelagic, the higher relative importance of *heterogeneous selection* in the Mediterranean Sea than in the open ocean is probably linked to its environmental gradients: north-south increasing temperature (45), west-east increasing salinity(45), and west-east decreasing nutrient concentrations (11). This result contradicts a previous hypothesis that *homogeneous selection* should be the most important process in all ocean basins (46). Instead, the balance between ecological processes shaping picoplankton communities will change depending on the analyzed environmental heterogeneity, circulation patterns, and geographic scale (47–49). In our study, the overall role of *selection* decreased, for both domains, when transiting from the epipelagic into the deep waters, where there is relatively lower environmental heterogeneity in comparison to the epipelagic (*SI Appendix*, Fig. S10). Moreover, the coupling between picoplankton community differentiation and environmental distances was stronger in the epipelagic than in the deep ocean, further indicating that the relative importance of selection raises with increasing environmental variability. *Selection* was also the most important process shaping picoplankton when ecological processes were estimated with all samples of our dataset, which captures environmental differences from surface to deep waters. This is another evidence that *selection* is enhanced as environmental heterogeneity is increased. These findings are coherent with ecological theory and other studies that show that high environmental heterogeneity leads to higher *selection (20)* in terrestrial (27, 50) and aquatic ecosystems (29, 51, 52). Conversely, the sum of *dispersal limitation* and *drift* were overall higher in the deep than in the surface ocean, suggesting that factors such as microbial abundances (i.e. low population sizes) (23) and physical barriers (strongly differentiated water masses and deep-sea bathymetry) (18) play an important role in the structuring of deep ocean picoplankton communities (Fig. 6). *Dispersal limitation* increased with depth probably because of decreasing turbulence (stable water masses and slow currents) (31) and the presence of straits and seamounts (53) that work as geographical barriers for microbial dispersal in the deep ocean (Fig. 6). Other studies have shown how strong physical barriers can limit microbial dispersal in soils (50), sediments (27), ponds (52) and, potentially, in the ocean (22).

**Figure 6.**
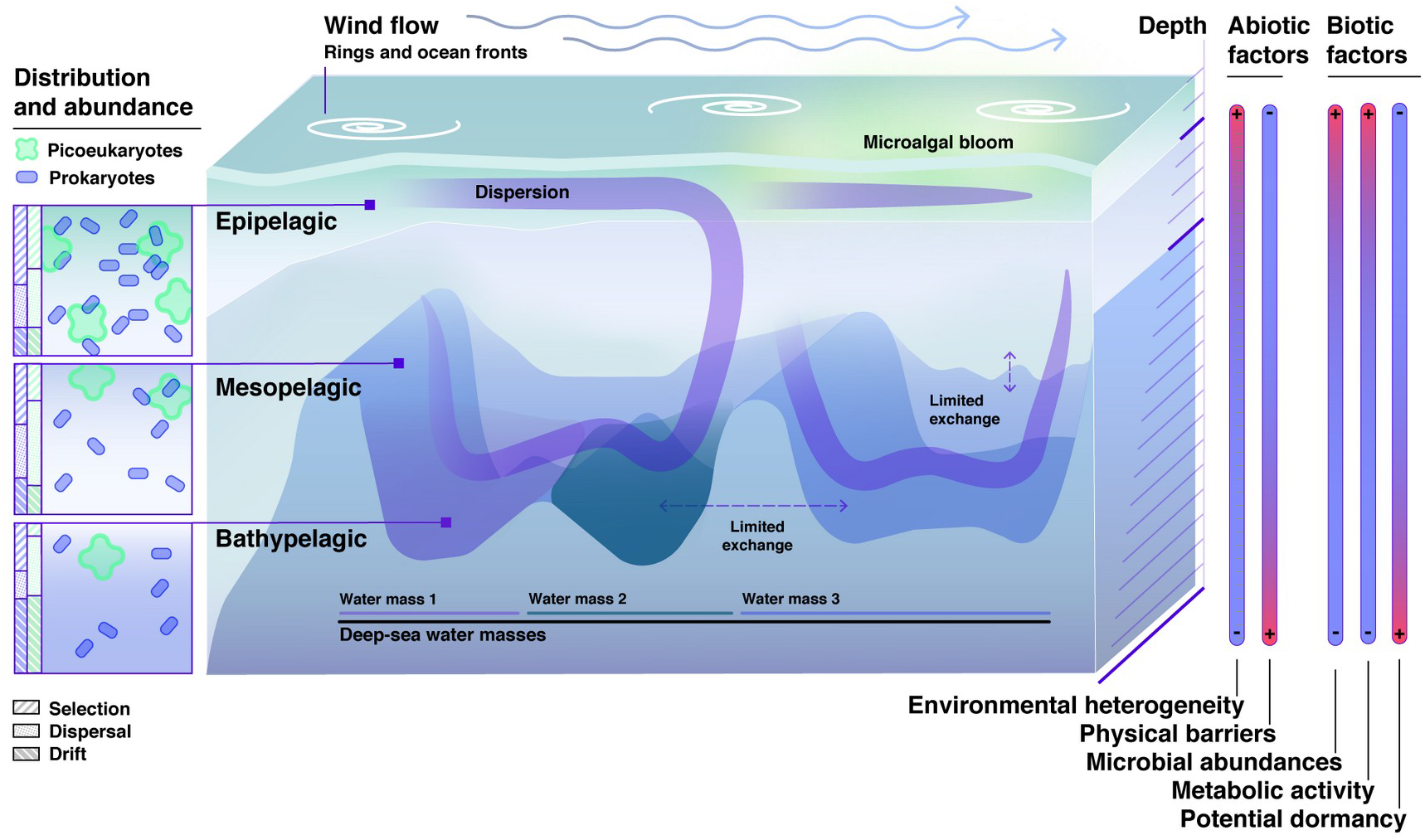
Conceptual model synthesizing the ecological processes assembling picoplankton communities across ocean depth zones. We used the main findings of this study and the knowledge available in the literature to construct this conceptual model. Vertical variation of biotic and abiotic factors, as well as geography (e.g., bathymetry), affect the ecological processes that generate community distribution patterns. The model predicts an increasing role of dispersal limitation with depth: *dispersal limitation* is weaker in the epipelagic than in the meso- and bathypelagic due to faster currents, and, potentially, aerial dispersal in surface waters, compared to more isolated deeper zones. Other mechanisms taking place in deep waters such as *a)* barriers to dispersal (e.g. water mass boundaries, deep sea topography) or *b)* limited random dispersal due to low species abundances, could also explain this pattern. *Selection* is the most important process structuring picoplankton communities in the epipelagic and displays a decreasing importance with depth due to higher habitat heterogeneity – driven by microalgal blooms, magnitude of the DCM and mesoscale processes (e.g.: ocean rings and fronts) – in upper than in bottom waters. The relative role of *drift* increases towards the deep, likely because of decreasing microbial abundances with depth. The importance of dispersal limitation is always higher in picoeukaryotes than in prokaryotes, given the smaller population sizes of picoeukaryotes and their limited capability to generate dormant stages to sustain long-range dispersal, compared to prokaryotes. Thus, a different balance of ecological processes assembles these domains, even when they share the same ocean zones.

### Water mass composition affects the distribution of prokaryotic communities

Water masses may impact microbial communities in basically two ways: *a)* as a selective force – since they have different temperatures and salinity (54, 55) as well as organic matter composition (56–58) – or; *b)* as a physical barrier to dispersal due to sometimes strong differences in water density (14). We found significant positive correlations between prokaryotic community structure and water mass compositions in the open ocean, which is in line with previous studies that found bacterial communities associated with specific water masses (13, 14, 56, 59). This relationship is likely linked to the fact that each water mass has different types of organic matter (57, 61) that likely select for different prokaryotes (56, 58). In turn, picoeukaryotes were only poorly correlated with differences in water mass in the open ocean, which implies that some of them could be able to swim across boundaries or that they are weakly linked to the composition of typical organic matter associated with each water mass. Instead, the high dispersal limitation of picoeukaryotes would be mainly regulated by their smaller population (9) as well as by their limited capability to enter into dormancy when compared to prokaryotes (62). In the Mediterranean Sea, the coupling between community and water mass composition was significant for both prokaryotes and picoeukaryotes in the meso- and bathypelagic, which agrees with previous reports (63) and it is likely linked to the strong horizontal cross-basin physical separation imposed by the Straits of Sicily and Gibraltar (11). Our results also point out that differences in both prokaryotes and picoeukaryotic communities are coupled with differences in water mass composition in vertical profiles (*SI Appendix*, Fig. S14). Interestingly, the slope and strength that differences in picoplankton composition was explained by differences in water masses varied among vertical profile stations (*SI Appendix*, Fig. S14). This result indicates that local-scale events (e.g. upwelling, dense water propagation) may also regulate the impact of water mass on microbial communities in a vertical dimension (64–66).

### Picoplankton communities display weaker biogeography in the surface than in the deep ocean

Our distance-decay analysis revealed that the autocorrelation in community and geographic distances is stronger in the deep than at the surface, which agrees with our sequential analysis results (Fig. 5B) and suggests that there are more marked changes across space in the deep ocean, particularly in the picoeukaryotic community. This result agrees with a recent study that found larger eukaryotic community dissimilarity between pairs of sites in the deep than in the surface global ocean (67). Such changes in community composition with increasing geographic distance (that is, distance-decay) can be generated by selection and/or dispersal limitation (68). For picoeukaryotes, the fact that changes in community composition were better explained by geography (ocean basin) than by environmental variation (Fig. 2B) supports that the distance-decay pattern in the deep sea is predominantly related to dispersal limitation (18, 67). On the other hand, prokaryotic community structure was predominantly explained by environmental variables rather than by geography (Fig. 2B) which indicates that, in this domain, distance-decay is mostly driven by selection. It is important to notice that, since many prokaryotes may be in dormant state (69), the distance-decay could have been stronger if we had analyzed measures of the active prokaryotic community (using RNA) instead of measures of the total community (with DNA), as previously shown for bacterial communities (69). Another important factor that could be increasing the role of *dispersal limitation* and *drift* in the deep ocean is the decreasing microbial population sizes from surface to deep waters. Rare species with small populations are less likely to disperse (70) and more likely to randomly collapse than species with large populations (23). As expected (9), microbial abundances drastically decreased towards the deep ocean so that the deep ocean contains only 1% of the organisms of the surface ocean (9). Overall, the depth-related patterns in ecological processes were more pronounced in the Mediterranean Sea than in the open ocean, which is partially explained by being a semi-enclosed sea, with unique oceanographic features such as limited circulation, sharp geographic barriers and strong environmental gradients (11, 40).

### Differences between picoplankton members in the different depth zones

A different balance of ecological processes shapes prokaryotic and picoeukaryotic communities in several ecosystems (34, 52, 71), including the surface ocean (17, 46). Here we found that such differences between domains persist in the deep ocean. *Dispersal limitation* was always higher for picoeukaryotes than for prokaryotes, which agrees with previous studies using similar approaches conducted in Antarctic lakes (51) and in basin-scale oceanic regions (49). This contrast between domains in terms of dispersal rates is partially due to organismal and population size differences (34, 70, 72). Unicellular eukaryotes are on average 3 times larger than prokaryotes and, therefore, would be expected to be more limited by dispersal (34, 72). Picoeukaryotes (∼10^3^ cells mL^-1^) have populations that are about three orders of magnitude smaller than prokaryotes (∼10^6^ cells mL^-1^), which decreases their likelihood to disperse (70). *Homogeneous selection* was in general higher in prokaryotes than in picoeukaryotes, which is in line with previous findings in the Pacific Ocean (46). This supports that environmental heterogeneity can act differently on prokaryotic and picoeukaryotic assembly across depths. The reason is likely due to different adaptations to the same environmental heterogeneity of prokaryotes and picoeukaryotes (62). For instance, a given degree of environmental heterogeneity could select for a few generalist species that have wide niches or many specialist species with narrow niches or a combination of both strategies. Moreover, the relatively higher *homogeneous selection* in prokaryotes than in picoeukaryotes suggests that dormancy could be playing an important role in modulating prokaryote assembly in the deep ocean. Dormancy is indeed a common mechanism in prokaryotes to overcome harsh environmental conditions (73). This mechanism has been shown to affect metacommunity structure by dampening distance-decay relationships and maintaining local diversity (69, 74, 75). Many prokaryotes reach the deep ocean from the surface through vertical dispersal (76) or disperse as endospores from sediments (77). However, DNA-based community composition data includes non-active bacterial cells (78), likely in dormancy state, to survive the very different conditions of the dark and cold deep ocean (77). Therefore, a relatively higher proportion of dormant bacteria can create an apparent ‘homogenization’ of prokaryotic communities in deep zones. In fact, evidence exists that bacteria decrease their activity towards the deep dark ocean [*SI Appendix*, Fig. S16] (9, 79). As far as we know, dormancy has not been reported in picoeukaryotes (62), which could partially explain the negligible role of *homogeneous selection* in the assembly of this domain in the deep ocean. Finally, we found that the higher spatial turnover (sequential horizontal changes) in picoeukaryotes than in prokaryotes in the surface ocean (17) is also observed in the deep ocean. Furthermore, we show that this difference in spatial turnover between domains increases with depth, which is coherent with dispersal limitation being an increasingly important processes shaping picoeukaryotic communities in deeper ocean zones.

### Potential picoplankton responses to multiple environmental changes across ocean depths

The global ocean is facing drastic changes in important environmental drivers such as temperature, pH, salinity, and nutrient concentrations (80, 81), which are very likely affecting all domains of life, their community structure, and interactions (82). Global climate change is also driving changes in ocean currents due to shifts in wind patterns, heat balance, and freshwater inflows from glacial melting (83), which may directly affect plankton dispersal rates (84). Water masses have also been modified by anthropogenic changes in temperature and salinity even in the deep ocean (85, 86), which may affect picoplankton community composition (13, 56, 60) by changing both selection and dispersal assembly processes. Our results suggest that the prokaryotic and eukaryotic components of the ocean’s smallest plankton are likely to respond differently to environmental change as a result of the different balance of ecological processes structuring their communities. Prokaryotes seem to be relatively more sensitive to selective forces than picoeukaryotes (17), so that changes in important environmental drivers (e.g. temperature, organic matter composition) will have a higher potential to affect prokaryotic community composition at a global scale (17, 58) than changes in dispersal drivers (e.g. currents, fronts). On the other hand, picoeukaryotic community composition at global scales would be potentially more affected by changes in factors regulating horizontal and vertical dispersal processes – such as current circulation (33) and thermal stratification (87) – than by environmental drivers. While here we refer to the entire community, specific picoeukaryotic taxa might be strongly structured by environmental drivers (88). Indeed, temperature is well-known to influence relatively more heterotrophic than photosynthetic eukaryotic activity (89). For instance, cosmopolitan unicellular picoeukaryotic predators (MAST-4) display clear temperature-driven niche-partitioning in the ocean (90). After all, in a long timescale, no matter the dispersal rate of a given species, it will eventually be selected and constrained by local abiotic and biotic factors (84). Thus, the relative effect of projected changes in environmental selection and dispersal pathways on microbial communities should be evaluated together.

Most importantly, our work suggests that the microbial communities inhabiting the deep ocean are likely to respond differently to environmental changes than those living in the surface ocean. This is particularly relevant in the context of increasing multiple stressors caused by climate change (warming, acidification, and deoxygenation) and human exploitation activities (i.e.: mining, oil and gas extraction, waste disposal) in the deep ocean (91). While upper ocean picoplankton communities would be relatively more sensitive to changes in environmental selective forces (e.g. temperature and nutrient concentration), deep ocean picoplankton communities should be relatively more impacted by the removal or creation of dispersal pathways. In this regard, projected perturbations in temperature, pH, oxygen, and nutrient concentration (80) should impact relatively more the small plankton communities inhabiting the upper than those in the deep ocean. Yet, changes in air fluxes and ocean currents should also affect the surface picoplankton community (92), but relatively less than selective forces. On the other hand, changes in dispersion vectors should be the main factor altering the balance of ecological processes assembling picoplankton communities in the deep ocean. For example, ocean micro- and nanoplastic pollution, a widespread environmental issue (93, 94) could represent important substrates for both prokaryotes and single-cell eukaryotes colonization and work as efficient dispersion vectors (95), potentially altering dispersal rates across ocean depth zones. Furthermore, changes in ocean stratification patterns are reducing nutrient exchange and expanding oligotrophic conditions in the upper ocean (96). Our vertical profile results suggest that this increased stratification could affect not only microbial selective forces, but also dispersal across depth zones. These changes can ultimately impact important ocean ecosystem services such as primary productivity and nutrient cycling at a global scale (87, 97, 98).

### A conceptual framework for the global biogeography of picoplankton across ocean depths

Historically, many studies have focused on the effect of *selection* – also referred to as niche-modeling or environmental filtering – on marine microbial communities (99–101). Other studies aimed to model how *dispersal* influences microbial biogeography in the global surface ocean (34, 102–104). More recently, there have been important efforts bringing together environmental selection and dispersal in the ocean (18, 33, 84). Nevertheless, besides selection and dispersal, picoplankton community assembly is also ruled by ecological drift (17, 29). Integrating these processes into a single framework considering organism, environmental and physical differences between depth zones was still missing. By combining empirical evidence, we propose a novel conceptual framework that expands the current understanding of plankton community assembly in environmentally distinct ocean depth zones (Fig. 6). It synthesizes how environmental heterogeneity, water mass structure, deep-sea topography, microbial abundance, and activity mediate the action of ecological processes assembling the two components of the smallest plankton communities (Fig. 6). This framework can be used to delineate hypothesis-driven studies to predict how plankton assemblages will respond across depths to multiple stressors in a changing ocean (105). For instance, based on this framework, we can expect that the balance between determinism (selection) and stochasticity (dispersal limitation or ecological drift) would decrease with plankton size. Thus, nano- (3-20 µm), micro- (20-200 µm), and mesoplankton (200-2,000 µm) biogeography would be increasingly limited by dispersal and display more marked biogeography (34, 88, 106), especially in the deep ocean (Fig. 6). We can also foresee that particle-attached prokaryotes – which are particularly relevant in the deep ocean (107) – should be more limited by dispersal than free-living prokaryotes. In general, the importance of dispersal limitation relative to that of selection should increase not only with organism and particle sizes, as expected by the size-dispersal hypothesis (108), but also with ocean depth. Here we show that this dispersal-selection balance, regulated by organism size, should be more pronounced in the deep than in the upper ocean.

## Methods

### Dataset, sampling, and analytical methods

We compiled a dataset (Fig. 1) composed of 451 samples from surface (3 m depth) to deep waters (up to 4,800 m), covering three depth zones of the ocean: epi- (0-200 m – including DCM), meso- (200-1,000 m), and bathypelagic (1,000-4,000 m). This dataset combines samples obtained during two oceanographic expeditions with similar sampling strategies: *i)* the *Malaspina-2010* circumglobal expedition (41, 109) from which we included 263 samples collected between December 2010 and July 2011 in 120 stations distributed along the tropical and subtropical portions (latitudes between 35° N and 40° S) of the Pacific, Atlantic, and Indian oceans (Fig. 1); and *ii)* the *HotMix* trans-Mediterranean cruise (11, 61) from which we considered 188 samples collected between April and May 2014 in 29 stations distributed along the whole Mediterranean Sea (from -5° W to 33° E) and the adjacent Northeast Atlantic Ocean (Fig. 1A). This dataset therefore allows the comparison of the tropical and subtropical ocean (samples hereafter called “open ocean”) to a semi-enclosed basin such as the Mediterranean Sea, which displays unique features such as higher temperature and salinity as well as lower nutrient concentration than the open ocean, particularly in the meso- and bathypelagic (Fig. 1B). The *Malaspina-2010* contains 13 stations where the whole vertical profile was sampled (VP stations in Fig. 1). A detailed vertical distribution of the samples is available in the Supplementary Material (Fig. S1). Due to the difference in the sampling size between depth zones, we also generated subsets with a standardized number of samples (n=39) evenly-distributed across space (Fig. S7 and Fig. S8).

This dataset comprises a contextual database with a total of 6 standardized environmental parameters (temperature, salinity, fluorescence, PO43−, NO3−, and SiO2) as well as prokaryote and picoeukaryote abundances determined by flow cytometry and bacterial activity measurements. Water samples were obtained with 20L (in *Malaspina*) or 12L (in *HotMix*) Niskin bottles attached to a rosette sampler equipped with a conductivity–temperature–depth (CTD) profiler (except surface samples in *Malaspina*, that were obtained with individual 30 L bottles, not attached to the rosette). Vertical profiles of temperature, conductivity, and fluorescence were continuously recorded throughout the water column with the CTD sensors. Conductivity measurements were converted into practical salinity scale values. Inorganic nutrients (NO_3_^−^, PO_4_^3−^, SiO_2_) were measured from the Niskin bottle samples with standard spectrophotometric protocols (110), using a Skalar autoanalyzer SAN++, as described in (41, 57). Missing nutrient concentration values were extracted from the World Ocean Database (111). Prokaryotic populations and phototrophic picoeukaryotes abundances were enumerated using a FACSCalibur flow cytometer (BD Biosciences, San Jose, CA, USA) as detailed elsewhere (112). Prokaryotic heterotrophic activity was estimated using the centrifugation method and measuring ^3^H-leucine incorporation (113). For deep water samples we used the filtration method with a larger volume and undiluted hot leucine. Significant differences in microbial abundances and bacterial activity between depth zones were tested with an analysis of variance (ANOVA), followed by a Tukey post-hoc test.

To obtain picoplankton biomass, ∼4–12 L of seawater were first pre-filtered with a 200-µm net mesh (to remove large organisms and particles). *Malaspina* samples were then sequentially filtered through a 20 μm nylon mesh followed by 3-μm and 0.2-μm polycarbonate filters (47-mm for surface and 142-mm diameter for vertical profiles, Isopore, Merck Millipore, Burlington, MA, USA) using a peristaltic pump. *HotMix* samples were sequentially filtered through 47-mm 3-μm policarbonate filters (Isopore, Merck Millipore) and 0.2-μm Sterivex units. Filters were flash-frozen in liquid N2 and stored at −80 °C until DNA extraction. Here, only the free-living ‘picoplankton’ size-fraction (0.2–3 μm) was used in downstream analyses.

### Nucleic acid extraction, sequencing, and bioinformatics

DNA extraction was conducted with a standard phenol-chloroform protocol (114) for the *Malaspina* surface samples. DNA from the *Malaspina* vertical profile samples was extracted using the Nucleospin RNAkit (Macherey-Nagel) plus the Nucleospin RNA/DNA Buffer Set (Macherey-Nagel) procedures. *HotMix* DNA samples were extracted using the PowerWater Sterivex™ DNA isolation Kit (MO BIO Laboratories). DNA extracts were quantified with Qubit 1.0 (Thermo Fisher Scientific) and preserved at −80 °C. The same extracts were used for both the 16S and 18S rRNA-gene amplification and all samples were sequenced with the same prokaryotic and eukaryotic primers. The hypervariable V4–V5 (≈ 400 bp) region of the 16S rRNA gene was PCR amplified with the primers 515F-Y (5’-GTGYCAGCMGCCGCGGTAA) -926R (5’-CCGYCAATTYMTTTRAGTTT) to target prokaryotes – both Bacteria and Archaea (115). The hypervariable V4 region of the 18S rRNA gene (≈ 380 bp) was PCR amplified with the primers TAReukFWD1 (5’-CCAGCASCYGCGGTAATTCC-3’) and TAReukREV3 (5’-ACTTTCGTTCTTGATYRA-3’) to target eukaryotes (116). PCR amplification was carried out with a QIAGEN HotStar Taq master mix (Qiagen Inc., Valencia, CA, USA). Amplicon libraries were then paired-end sequenced on an Illumina (San Diego, CA, USA) MiSeq platform (2 × 250 bp or 2 × 300 bp) at the Research and Testing Laboratory facility, Texas, USA (https://rtlgenomics.com/). See details about gene amplification and sequencing in (11, 17).

Raw Illumina miSeq reads (2×250 or 2×300) were processed using DADA2 (117) to determine amplicon sequence variants (ASVs). For the 16S rRNA gene, forward reads were trimmed at 220 bp and reverse reads at 200 bp, whilst for the 18S rRNA gene, we trimmed the forward reads at 240 bp and the reverse reads at 180 bp. Then, for the 16S, the maximum number of expected errors (maxEE) was set to 2 for the forward reads and to 4 for the reverse reads, while for the 18S, the maxEE was set to 7 and 8 for the forward and reverse reads respectively. Error rates for each possible nucleotide substitution type were estimated using a machine learning approach implemented in DADA2 for both the 16S and 18S. Unsurprisingly, error rates increased with decreasing quality score. Finally, DADA2 was used to estimate error rates for both the 16S and 18S genes in order to delineate the ASVs

Prokaryotic ASVs were assigned taxonomy using the naïve Bayesian classifier method (118) alongside the SILVA v.132 database (119) as implemented in DADA2, while Eukaryotic ASVs were BLASTed (120) against the Protist Ribosomal Reference database [PR^2^, version 4.11.1; (121)]. Eukaryotes, chloroplasts, and mitochondria were removed from the 16S ASVs table, while Streptophyta, Metazoa, and nucleomorphs were removed from the 18S ASVs table. Both, the 16S and 18S ASVs tables were rarefied to 20,000 reads per sample with the function *rrarefy* from the Vegan R package. To be consistent with our previous study (17), for the calculation of ecological processes and associated analysis, ASVs with total abundances < 100 reads across all samples were removed to avoid PCR and sequencing depth biases. This filtering procedure removed ∼5% of the total reads and ∼90% of the total ASVs from both the 16S and the 18S rRNA datasets.

Computing analyses were conducted at both the MARBITS bioinformatics platform of the Institut de Ciències del Mar (ICM; http://marbits.icm.csic.es) and the MareNostrum (Barcelona Supercomputing Center). Sequences are publicly available at the European Nucleotide Archive (http://www.ebi.ac.uk/ena) under accession numbers PRJEB23913 [18S rRNA genes] & PRJEB25224 [16S rRNA genes] for the *Malaspina* expedition; PRJEB23771 [18S rRNA genes] & PRJEB45015 [16S rRNA genes] for the *Malaspina* vertical profiles; and PRJEB44683 [18S rRNA genes] & PRJEB44474 [16S rRNA genes] for the *HotMix* expedition.

### Phylogenetics

Phylogenetic trees were built for both the 16S and 18S rRNA gene-datasets using the ASVs full sequences. Raw ASV sequences were firstly aligned against an aligned SILVA template – for 16S rRNA – and an aligned PR^2^ template – for 18S rRNA – using mothur (122). Poorly aligned regions or sequences were then removed using trimAl (parameters: -gt 0.3 -st 0.001) (123). Aligned sequences were also visually curated with seaview v4 (124) and sequences with >=40% of gaps were removed. Finally, phylogenetic trees were inferred from the aligned quality-filtered sequences using FastTree v2.1.9 (125). Additional phylogenetic analyses were carried out with the *picante* R package (126).

### Environmental heterogeneity, water masses characterization, and least-cost distance calculations

We calculated the average pairwise dissimilarity (*EnvHt*) as an index of environmental heterogeneity based on the main standardized environmental variables: temperature, salinity, fluorescence, PO_4_^3−^, NO_3_^−^, and SiO_2_. We firstly computed an Euclidean distance matrix for each depth zone using the *vegan* R package and then determined the dissimilarity among samples by dividing the Euclidean distance matrix (*Euc*) by the maximum Euclidean distance (*Eucmax*) of a given depth zone as described in (29) and summarized here: *EnvHt*=(*Euc*/*Eucmax*)+0.001. Finally, the mean *EnvHt* (*EnvHt*) was calculated as an estimation of environmental heterogeneity in each depth zone. Significant differences in environmental heterogeneity between depth zones were tested with a Kruskal-Wallis test, followed by a Wicoxon post-hoc test.

The presence of different water masses is an important feature to properly describe the deep dark ocean ecosystem (> 200 m depth). Water masses are well-established water bodies with unique properties that can be characterized by their thermohaline and chemical features. A water mass is composed of different proportions of one or more water types of a given origin (127). Here, the percentage of different water types contributing to the water mass composition of each sample (from 200 m to the bottom) was calculated using an optimum multiparameter water mass analysis (128). This method basically characterizes water types by using conservative variables such as salinity and potential temperature (see (61) for details). We have identified 22 and 19 water types in the open ocean and in the Mediterranean Sea, respectively. We computed the dissimilarity (Euclidean distance) between pairwise samples based on their water mass composition (% of each water type) to use in our downstream analysis. A nonmetric multidimensional scaling (NMDS) analysis based on these euclidean distances was conducted to determine the differences among samples.

Least-cost geographical distances were calculated using the ‘lc.dist()’ function of the *marmap* R package (129). We first computed three transition matrices (using the ‘trans.mat()’ function) with different minimum depths, corresponding to the epi- (surface), meso- (200 m), and bathypelagic (1,000 m). Each generated transition matrix contained the probability of transition from one cell to adjacent cells of a given bathymetric grid. We used the high-resolution (15 arc-second) GEBCO bathymetric database hosted on the British Oceanographic Data Centre server (https://www.gebco.net/). Since the Mediterranean Sea deep waters (>400 m) are completely separated by the Strait of Sicily, the *marmap* algorithm could not calculate the horizontal distance between bathypelagic samples situated in the western and eastern Mediterranean. To deal with this issue, we simulated the vertical trajectory needed to overcome the Strait of Sicily by simply summing each sample’s depth to the geographical distances between ‘isolated’ stations. To calculate the least-cost distances, ‘marmap’ sets a depth limit for geographic barriers to compute the transition matrices (129). For example, if the limit is set to 0, the program calculates the distance turning around the continents. However, in the case of the Mediterranean Sea, the western and eastern basins are completely isolated (at least horizontally) in depths down to 400m, so the program outputs unrealistic very long distances between western and eastern samples from the deep ocean. To deal with this issue, for these isolated samples, we computed the least-cost distances by calculating the normal geographic distances (geodesic) between samples (not considering geographic barriers) and then summed the vertical distances to theoretically overcome the Strait of Sicily. For example, a western 1,400 m depth sample (1 km deeper than the top of the Strait of Sicily) located 200 km from an eastern 1,400 m depth sample had a final least-cost distance of 200 km + 2x 1 km = 202 km.

### Quantification of the ecological processes

The action of ecological processes (selection, dispersal, and drift) were here quantified using a null model approach (27) that has been successfully applied to microbial ecology studies in diverse aquatic environments (29, 51, 52, 130). This analysis consists of two main sequential steps: *1)* inference of *selection* from ASV phylogenetic turnover; and *2)* inference of *dispersal* and *drift* from ASV compositional turnover (27). Since the existence of a phylogenetic signal (131) is an assumption of the first step of this method (27), we first tested whether closely related taxa (based on the 16S and 18S rRNA-gene phylogeny) were more similar in terms of habitat preferences than distantly related taxa. Mantel correlograms between ASVs niche and phylogenetic distances were used to test for a phylogenetic signal in the variables that explained the highest fraction of community variance in each depth zone. We detected a phylogenetic signal within short phylogenetic distances, which is in line with the literature (17, 27, 29).

Having fulfilled this assumption, we determined the phylogenetic turnover using the abundance-weighted β-mean nearest taxon distance (βMNTD) metric (27), which computes the mean phylogenetic distances between each ASV and its closest relative in each pair of communities (pairwise comparisons). Afterward, we run null models with 999 randomizations to simulate the community turnover by chance (βMNTDnull), in other words, without *selection* influence (27). Finally, the β-Nearest Taxon Index (βNTI) was calculated from the differences between the observed βMNTD and the mean βMNTDnull values. Overall, |βNTI| > 2 indicates that taxa are phylogenetically more related or less related than expected by chance, pointing to a strong influence of selection on community assembly (27). More precisely, βNTI values higher than +2 indicate the action of heterogeneous selection, while βNTI values lower than –2 points out to the action of homogeneous selection (27).

The β-diversity of communities that were not governed by selection (|βNTI| ≤ 2) was evaluated in a second step, which consisted of computing ASV taxonomic turnover to calculate the influence of either dispersal or ecological drift on community structure. To do so, we calculated the Raup-Crick metric (132) based on the Bray-Curtis dissimilarities (RCbray) (27). RCbray compares the measured β-diversity against the β-diversity obtained from null models (999 randomizations), representing a random community assembly (ecological drift). Absolute RCbray values smaller than (|RCbray| ≤ 0.95) indicate a community assembled by ecological drift alone (i.e., by chance). On the other hand, RCbray values > + 0.95 or < − 0.95 indicate that community assembly is structured by dispersal limitation or homogenizing dispersal, respectively (132). To further investigate the community assembly patterns in each depth zone, we used the ‘betapart’ R package (133) to calculate the partitioning of β-diversity (Jaccard, Sorensen and Bray-Curtis) into turnover or nestedness (134).

The relative importance of ecological processes were calculated for each depth zone subset. Additionally, we globally calculated these processes by integrating all depths of both datasets (Fig. S5). Since there are processes taking place along the water column (vertically) that may impact the biogeography that we observe horizontally in each depth zone, we also estimated the ecological processes integrating all depths (from 3 to 4,000 m) in each of the 13 vertical profile stations (Fig. 1A; see also Fig. S1 for sample vertical distribution).

### General analysis

Distance-based redundancy analyses (dbRDA) were performed on community composition (based on Bray-Curtis dissimilarities) of both prokaryotic (16S rRNA gene) and picoeukaryotic (18S rRNA gene) samples using the ‘capscale()’ function of the *vegan* R package (135). Analyses of dissimilarities were conducted using the ‘adonis2()’ function of the *vegan* R package to investigate the percentage of variance in community composition explained by environmental or geographic variables (136). Classic biogeographic provinces classifications (e.g.: Longhurst provinces; (137)) are only applied to the upper sunlit ocean (above 200 m), while deep-oceanic basins classifications (based on isolated water bodies) are only applied to the deep (bellow 3,500 m) (35). Therefore, we here used the classic geographic oceanic basins (South Atlantic Ocean, North Atlantic Ocean, North Pacific Ocean, South Pacific Ocean and Indian Ocean) as a standard categorical explanatory variable to compare the effect of geography between depth zones of the open ocean. For the Mediterranean Sea, we used the sub-basin classification (Levantine Sea, Ionan Sea, Sicily Strait, Tirrenyan Sea, Sardinian Sea, Alborean Sea and Gibraltar Strait), based on Mediterranean internal circulation patterns (138) as well as physico-chemical and biological features (139).

Spearman correlations were computed between β-diversity (bray-curtis and βNTI) and environmental euclidean distances matrices using the ‘cor.test()’ function of the *stats* R package. Spearman correlations were also carried out to test the association between community (bray-curtis dissimilarity) and water masses composition (euclidean distances) in the meso- and bathypelagic. Mantel correlograms were carried out with the ‘mantel.correlog()’ function in *Vegan* to test for the decrease in picoplankton community similarity (β-diversity) with increasing geographic distances (distance-decay). For the open ocean, we used distance classes of 1,000 km, while for the Mediterranean Sea we used distance classes of 350 km. Sequential differences in picoplankton β-diversity (bray-curtis dissimilarity) were computed in the sampling order of each project (see arrow directions in Fig. S15). Statistical differences between zones in sequential bray-curtis values were tested using analysis of variance (ANOVA) followed by a Tukey post-hoc test.

Pearson correlation matrices between diversity metrics and environmental variables were computed using the ‘cor()’ function and plotted with the *ggcorrplot* R package. Nonmetric multidimensional scaling (NMDS) based on Euclidean distances was used to visualize clustering in water mass composition among ocean depth zones and basins, followed by an analysis of similarities (ANOSIM) to test for differences among groups. The NMDS and ANOSIM were completed using the ‘metaMDS()’ and ‘anosim()’ *vegan* functions, respectively. Analysis of variance (ANOVA), followed by a Tukey post-hoc test, was used to test statistical differences in β-diversity metrics (Bray-Curtis, βNTI and RCbray). Differences in environmental heterogeneity values between zones were tested using Kruskal-Wallis, followed by a Wicoxon post-hoc test. Linear regression models were carried out to investigate the influence of water masses (euclidean distance) on community composition (bray-curtis dissimilarity) in each vertical profile. Spearman correlation was used to test correlation between the ecological processes results obtained with the total (unbalanced) dataset and the results found with a standardized sampling size dataset. All statistical analyses were conducted in the R statistical environment (140) and all plots were generated using the R package *ggplot2* (141).

## Data availability and resources

DNA sequences and environmental metadata are publicly available at the European Nucleotide Archive (http://www.ebi.ac.uk/ena) under accession numbers PRJEB23913 [18S rRNA genes] & PRJEB25224 [16S rRNA genes] for the *Malaspina* expedition; PRJEB23771 [18S rRNA genes] & PRJEB45015 [16S rRNA genes] for the *Malaspina* vertical profiles; and PRJEB44683 [18S rRNA genes] & PRJEB44474 [16S rRNA genes] for the *HotMix* expedition. R-Scripts for calculating the β-NTI and the Raup-Crick metrics are available at https://github.com/stegen/Stegen_etal_ISME_2013. The r-scripts used to generate figures and statistical analysis are available at: https://github.com/pcjunger/EcoProc_OceanDepths

## Acknowledgments

We are grateful to all scientists and crews from *Malaspina*-2010 and *HotMix* expeditions. Bioinformatics analyses were performed at the MARBITS platform of the Institut de Ciències del Mar (ICM http://marbits.icm.csic.es). PCJ was supported by Fundação de Amparo à Pesquisa do Estado de São Paulo – FAPESP (PhD grants #2017/26786-1 and #2020/02517-4) and by FAI/UFSCar (ProEx nº 3213/2020-83) through the European Union – H2020 project AtlantECO (award n° 862923). HS gratefully acknowledges continuous funding through Research Productivity Grants provided by CNPq (Process: 303906/2021-9). This work was supported by the projects INTERACTOMICS (CTM2015-69936-P, MINECO, Spain), MicroEcoSystems (240904, RCN, Norway), and MINIME (PID2019-105775RB-I00, AEI, Spain) to RL, and PID2021-125469NB-C31 to JMG, and by project HOTMIX (CTM2011-30010-C02-01 and CTM2011-30010-C02-02) of the Spanish Ministry of Economy and Innovation, co-financed with FEDER funds, to JA and JMG, respectively. We thank X. Antón Álvarez-Salgado for conducting the optimum multiparameter water mass analysis. The authors also thank Victor Saito, Melina Devecelli, Paula Huber and Cèlia Marrasé for their critical reading of an earlier version of this manuscript. The ICM authors acknowledge the *‘Severo Ochoa Centre of Excellence’ accreditation (CEX2019-000928-S)* to the ICM-CSIC.

