## Supplementary Material for "Global biogeography of the smallest plankton across ocean depths"

**This PDF file includes:**

SI Results
SI Discussion
SI References
Figures S1 to S16

**SI RESULTS**

In general, we observed an inverted diversity pattern between the two main components of the picoplankton community: while prokaryotic diversity (richness, Shannon index, and phylogenetic diversity) increased with depth, picoeukaryotic diversity decreased towards the deep ocean (Fig. S2). The Pielou evenness index increased for prokaryotes and decreased for picoeukaryotes from the surface to the deep ocean (Fig. S2). The gamma diversity of prokaryotes was 26776 ASVs in the open ocean and 11795 ASVs in the Mediterranean Sea. The gamma of picoeukaryotes was 35165 ASVs in the open ocean and 12367 ASVs in the Mediterranean Sea. The prokaryotic gamma diversity (adjusted to a standard sampling size; n=38) increased from the epi- (4203 in the open ocean and 2723 in the Mediterranean Sea) to the meso- (8898 in the open ocean and 4303 in the Mediterranean Sea) and bathypelagic (8898 in the open ocean and 4303 in the Mediterranean Sea). The picoeukaryotic adjusted gamma diversity (n=38) decreased from the epi- (12290 in the open ocean and 5699 in the Mediterranean Sea) to the meso- (6496 in the open ocean and 4217 in the Mediterranean Sea) and bathypelagic (4668 in the open ocean and 2330 in the Mediterranean Sea).

Prokaryotes and picoeukaryotes displayed significant differences ($p<0.05$) between depth zones for bNTI and $RC_{Bray}$ metrics (Fig S3). We also found significant differences ($p<0.05$) between ocean zones with regards to additional β-diversity metrics (i.e. Bray-Curtis, Jaccard and Sorensen) and their partitioning (Fig. S4). In the surface open ocean (~3m), the role of selection was higher for prokaryotic (~27%) than picoeukaryotic communities turnover (~11%) (Fig. S6). *Heterogeneous selection* had a relatively higher importance in structuring picoeukaryotes as compared to prokaryotes (~7% vs. ~4%, respectively). Conversely, *homogeneous selection* was higher for prokaryotes (~23%) than picoeukaryotes (~4%) (Fig. S6). *Dispersal limitation* explained ~67% of picoeukaryotic communities and ~25% of prokaryotic turnover in the surface open ocean (~3m). *Drift* was a relevant process structuring prokaryotes (~31%), but not for picoeukaryotes (~6%) (Fig. S6). Similarly, *selection* explained ~26% of prokaryotes and ~10% of picoeukaryotes turnover, in the surface Mediterranean Sea (Fig. S6). Unlike the open ocean, *heterogeneous selection* (~17%) explained a higher percentage of prokaryotes pairwise comparisons than *homogeneous selection* (~9%). An equal proportion of *heterogeneous* (~5%) and *homogeneous selection* (~5%) explained the structure of picoeukaryotes in the surface Mediterranean (Fig. S6). *Dispersal limitation* was the most important process (~63%) shaping picoeukaryotes and explained ~23% of the turnover of prokaryotes (Fig. S6), whereas *drift* explained most of the turnover of prokaryotes (~46%) and ~25% of picoeukaryotic assembly in the Mediterranean's surface waters (Fig. S6).

In the open ocean DCM, *selection* explained ~21% and ~46% of the turnover of prokaryotes and picoeukaryotes, respectively(Fig. S6). While *heterogeneous selection* was relatively more important for picoeukaryotes (~45%) than prokaryotes (~10%), *homogeneous selection* was much smaller for picoeukaryotes (0.8%) than prokaryotes (11%) in the DCM of the open ocean (Fig. S6). *Dispersal limitation* was relatively more important for prokaryotes (52%) than picoeukaryotes (37%) in this zone. *Homogenizing dispersal* was very low for both prokaryotes (~4%) and picoeukaryotes (~0.6%) in the DCM. *Drift* explained 26% and 13% of the turnover of prokaryotes and picoeukaryotes, respectively, in the DCM of the open ocean. In the Mediterranean Sea, *selection* was also relatively more important for picoeukaryotes (~23%) than prokaryotes (~16%) assembly (Fig. S6). *Heterogeneous selection* explained a larger proportion of the turnover of picoeukaryotes (~22%) than prokaryotes (~9%), whereas *homogeneous selection* was a more important process structuring prokaryotes (~7%) than picoeukaryotes (~0.4%) in the DCM of the Mediterranean Sea (Fig. S6).

Since there is also vertical dispersal between ocean surface and deep waters, we have also estimated the ecological processes integrating all depths (from 3 to 4,000 m) in each of the 13 vertical profile stations (VP stations in Fig. 1A). We found that *selection* was consistently the most important process vertically shaping free-living picoplankton communities in the *Malaspina* vertical profiles, explaining ~52-81% of the prokaryotic community turnover (Fig. S9) and ~24-52% of the picoeukaryotic community turnover (Fig. S9). The role of vertical *heterogeneous selection* ranged from 29% to 52% in prokaryotes and from 10% to 52% in picoeukaryotes (Fig. S9). *Homogeneous selection* was much higher for prokaryotes than picoeukaryotes in most of the *MalaVP* stations (Fig. S9). The role of vertical *dispersal limitation* ranged from 10% to 43% in prokaryotes and from 5% to 43% in picoeukaryotes (Fig. S9). Vertical *homogenizing dispersal* was negligible (~0-5%) for both domains (Fig. S9). The role of *drift* was relatively more important in picoeukaryotes (~15-43%) than in prokaryotes (~5-24%) across vertical profiles (Fig. S9).

Environmental heterogeneity was significantly higher in the epipelagic than in the meso- and bathypelagic of the open ocean and the Mediterranean Sea (Fig. S10). The bathypelagic displayed slightly higher environmental heterogeneity than the mesopelagic in the open ocean (Fig. S10). Instead, the mean environmental heterogeneity was the lowest in the Mediterranean bathypelagic (Fig. S10). Water masses composition was vertically structured in the open ocean (stress = 0.19521) with two separated clusters, meso- and bathypelagic, along the NDMS1 axis (Fig. S13). There was also a secondary separation between the Atlantic ocean and the Pacific+Indian oceans along the NMDS2 axis (Fig. S13). The ANOSIM analysis confirmed that this separation in water masses composition (euclidean distance) between zones (r=0.48, p<0.05) was stronger than that between ocean basins (r=0.39, p<0.05). In the Mediterranean Sea, there was a strong (stress = 0.1828) horizontal segregation in water masses between the Western and Eastern Mediterranean basins along the NMDS1 axis (Fig. S13). This segregation between basins was much stronger (r=0.52, p<0.05) than between zones (r=0.18, p<0.05) in the Mediterranean Sea.

### SI DISCUSSION

Both microbial domains (prokaryotes and picoeukaryotes) of the smallest ocean plankton communities are strongly structured by depth, which is overall in line with previous reports (1–4). Our results also endorse that the two domains of the picoplankton community have an inverted diversity pattern: prokaryotic diversity increased, whereas picoeukaryotic diversity decreased with depth. This pattern agrees with other studies performed in the Mediterranean Sea (1, 5). and in the open ocean at basin's (6–8) and global scales but covering only until 1,000 m depth (3). Here we confirm this depth diversity pattern – for both domains together – using a unique dataset covering the entire water column (down to 4,500 m depth) of the global tropical and sub-tropical ocean, including the Mediterranean Sea.

The relatively higher importance of *selection* that we found in the epipelagic (0-200m) slightly differs from the results that we found in the surface ocean (only ~3 m depth), where *dispersal limitation* and *drift* were more important than *selection* (9). This difference is probably because here we calculated the ecological processes in the epipelagic (0-200 m) using not only the surface, but also the DCM zone (Fig. S6), which can enhance environmental heterogeneity in the epipelagic (10) and lead to a different picoplankton composition in comparison to the SRF (3). Although the deep ocean is environmentally more homogeneous than the epipelagic, selection also played a relevant role in the deep. This finding is not surprising when we consider that the surface may influence the environmental variability in the deep ocean through dense water formation (11, 12), organic carbon fluxes (13) or even animal vertical migration (14). Besides, differences in water masses may confer some environmental heterogeneity in the deep ocean.

Overall, our framework represents a conceptual improvement in our understanding of microbial plankton community assembly in the global ocean. However, we recognize there are important eco-evolutionary mechanisms that were not included due to methodological constraints. For example, how does the role of microbial dormancy in community assembly change across ocean zones? Our framework suggests that it should increase with depth, but actual measurements of dormancy are required to confirm this prediction. How does microbial diversification (15) change across ocean depth zones? Are surface microbial communities more affected by diversification due to their higher metabolic and replication rates (16) as well as the faster environmental changes in the upper than in the deep ocean (17)? Or does the high dispersal limitation promote diversification (18) in the deep ocean? How do changes in these ecological processes couple with essential functions performed by the tiniest ocean plankton (19, 20)? These are some of the remaining questions to be addressed in future studies on ocean plankton biogeography.

### SI FIGURES

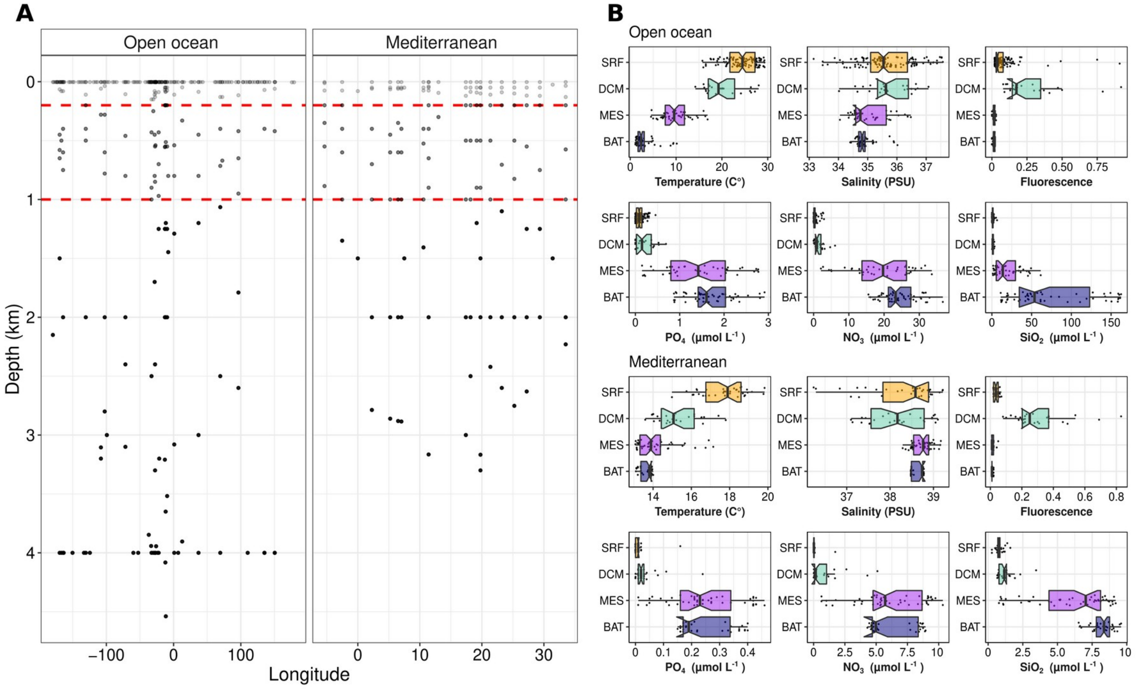


**Figure S1. (A)** Distribution of the samples across depth and longitude. The dashed red lines depict the division between zones: epi- (0-200 m), meso- (200-1,000 m) and bathypelagic (>1,000 m) **(B)** Boxplots showing the data variability, by depth zones, of the environmental variables used in this study. Note the difference in scales between the open ocean and the Mediterranean Sea. Means were significantly different (ANOVA, Tukey post-hoc test; $p<0.001$) between upper (SRF and DCM) and deep (MES and BAT) zones.

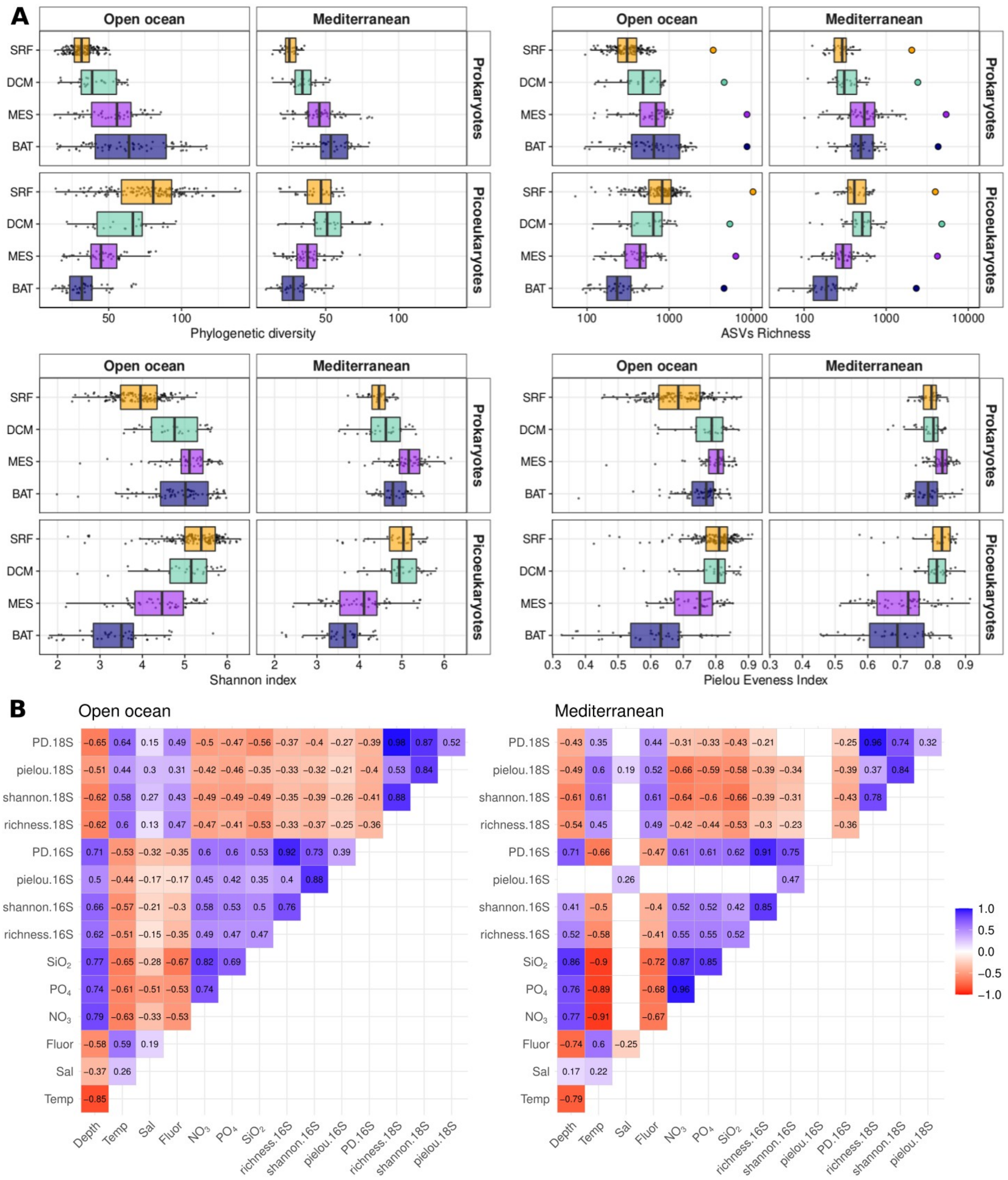


**Figure S2. (A)** Picoplankton diversity depicted as phylogenetic diversity, ASVs richness, shannon and Pielou evenness index by depth zones (SRF, surface; DCM, deep chlorophyll maxima; MES, Mesopelagic; BAT, Bathypelagic). The circles in the ASVs richness boxplots stand for gamma diversity adjusted by sampling size. **(B)** Correlation matrix of *Pearson* (R) correlation values between picoplankton diversity metrics and environmental variables in the open ocean and the Mediterranean Sea. The empty boxes represent non-significant correlations ($p$>0.05). The ‘16S’ tags in the metrics depict prokaryotic communities, while ‘18S’ tags represent picoeukaryotic communities. PD = phylogenetic diversity; Temp = Temperature; Sal = Salinity; Fluor = Fluorescence.

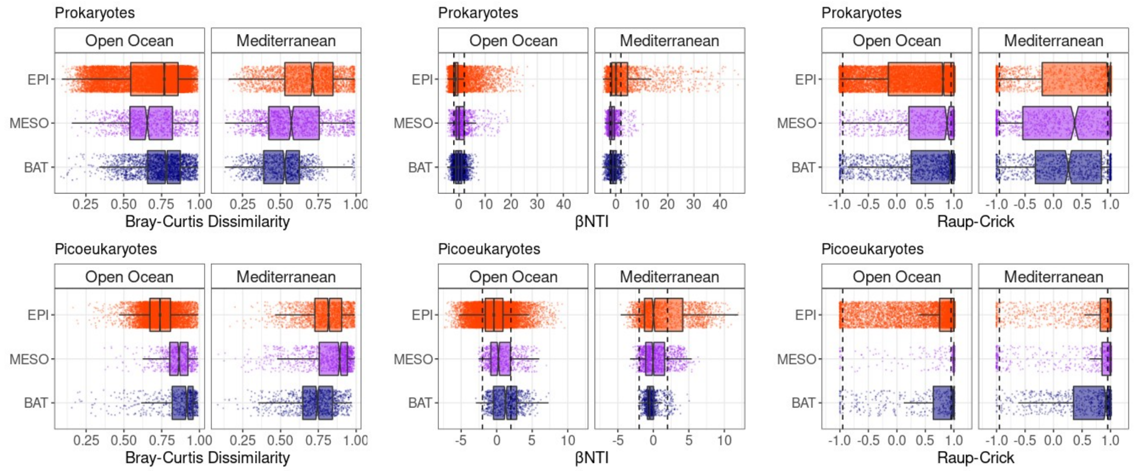


**Figure S3. Bray-Curtis, βNTI and $RC_{bray}$ metrics by depth zones** for **(A)** prokaryotes and **(B)** picoeukaryotes. See Fig. S4. for β-diversity partitioning plots. Means were significantly different (ANOVA, Tukey post-hoc test; p<0.001) between depth zones for both prokaryotes and picoeukaryotes.

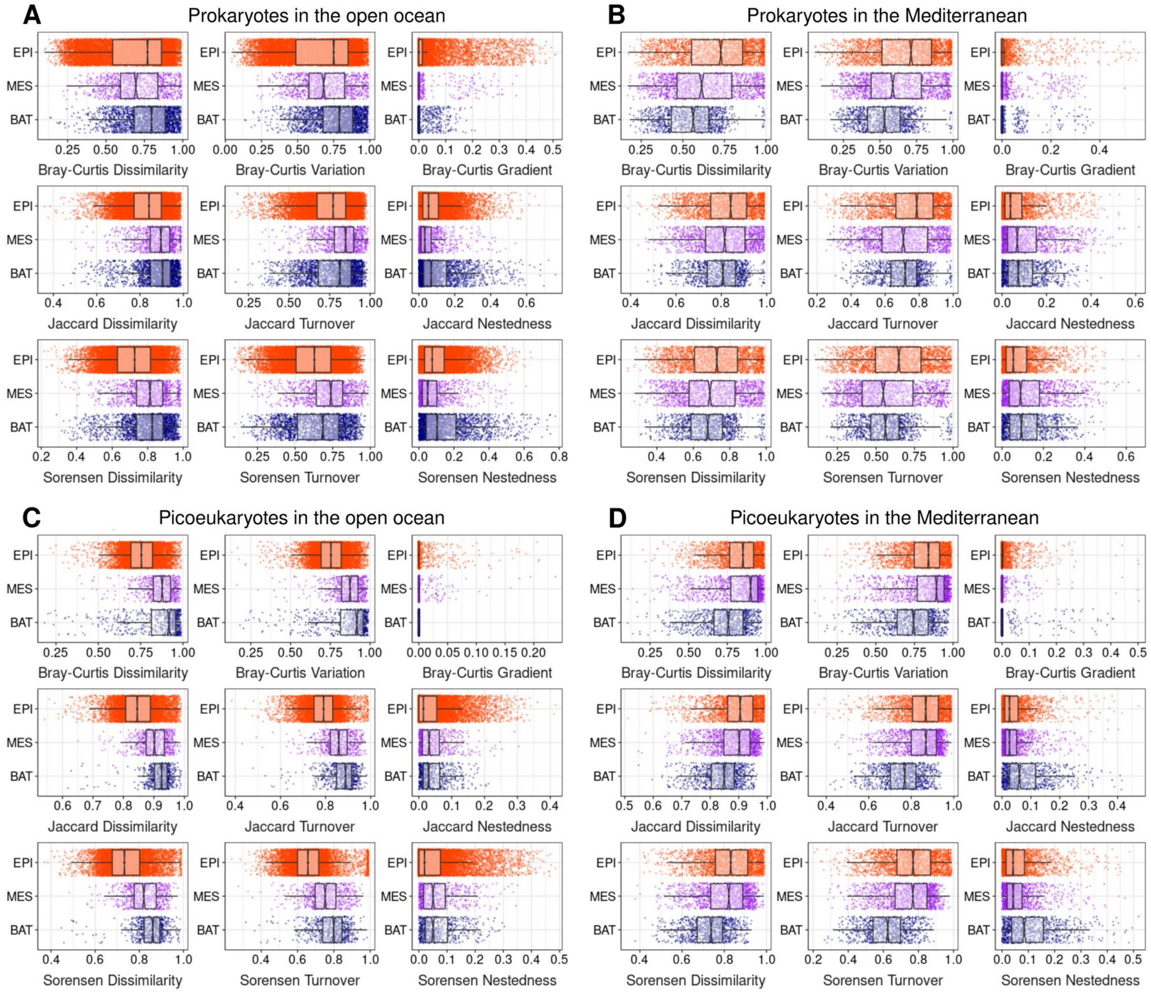


**Figure S4. Picoplankton β-diversity partitioning in the different ocean depth zones.** Bray-Curtis dissimilarity (variation and gradient), Jaccard dissimilarity (turnover and nestedness) and Sorensen (turnover and nestedness) for prokaryotes and picoeukaryotes in the open ocean (left panels) and Mediterranean Sea (right panels). Means were significantly different (ANOVA, Tukey post-hoc test; p<0.001) between depth zones for both prokaryotes and picoeukaryotes.

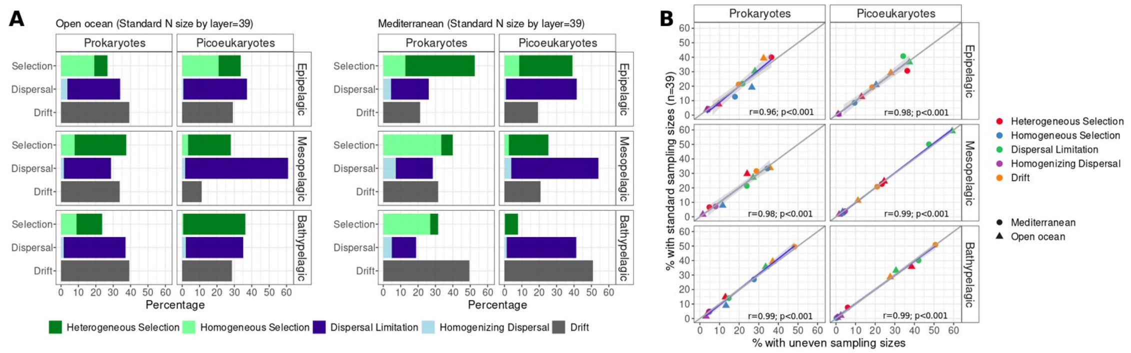


**Figure S5. Picoplankton community assembly processes across ocean depth zones using standardized sampling sizes (n=39). (A)** Relative importance of the ecological processes structuring the picoplankton communities at different depth zones of the open ocean and Mediterranean Sea: Epi- (n=39), Meso- (n=39) and Bathypelagic (n=39). **(B)** Linear regression between results obtained with total (unbalanced) and standardized sampling size dataset. These samples were evenly distributed across space as shown in Fig S6.

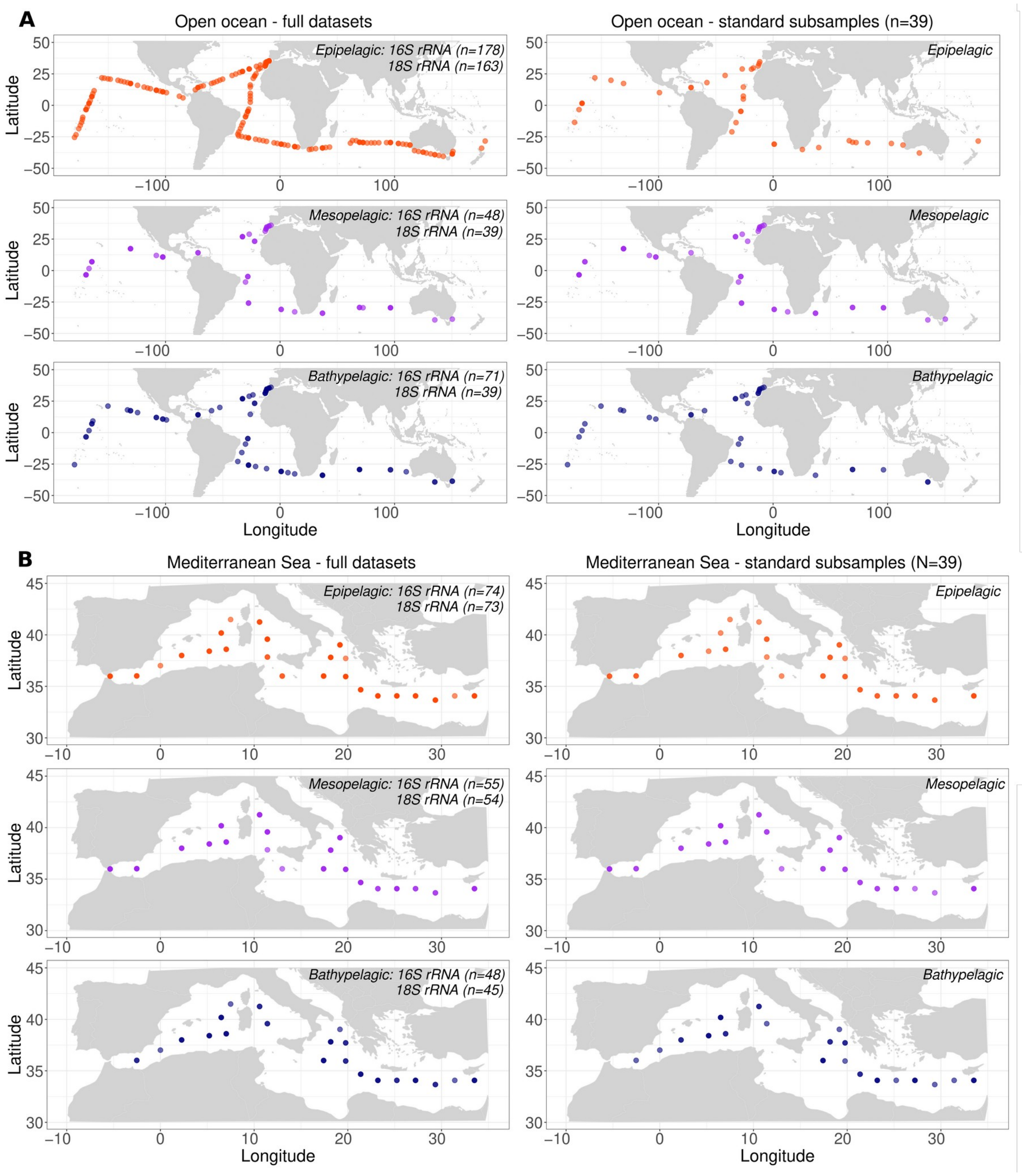


**Figure S6.** Geographic distribution of sampling stations in each zone of full datasets and subsets with standardized sampling size (n=39) in the **(A)** open ocean and **(B)** Mediterranean Sea. Samples were evenly distributed across space.

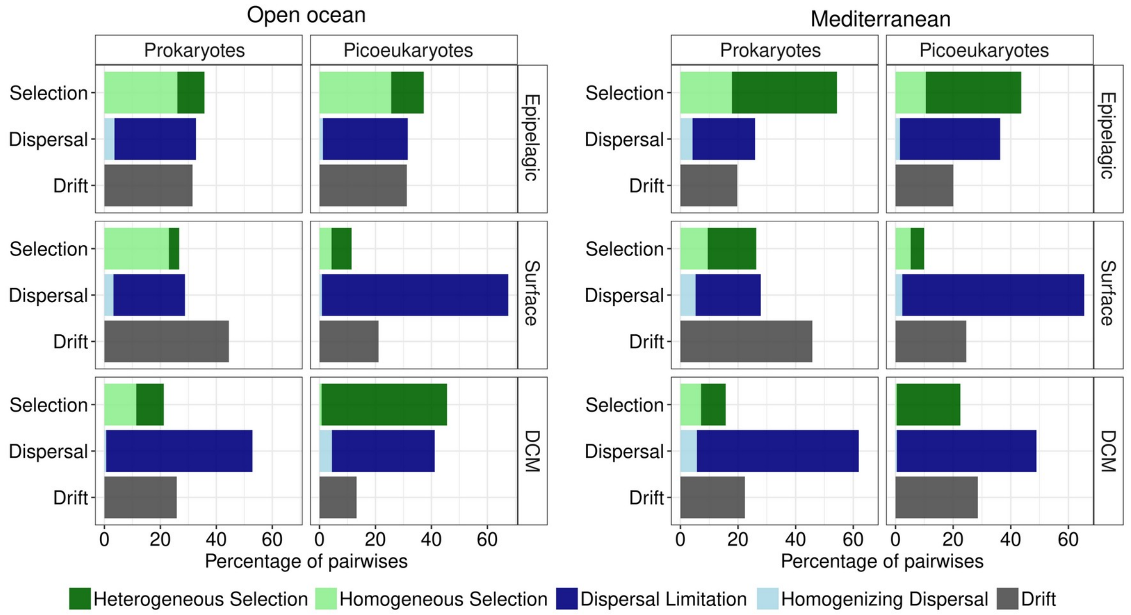


**Figure S7. Picoplankton community assembly processes in distinct deph zones of the epipelagic.** Relative importance of the ecological processes structuring the picoplanktonic community in the SRF and DCM zones of the open ocean and Mediterranean Sea.

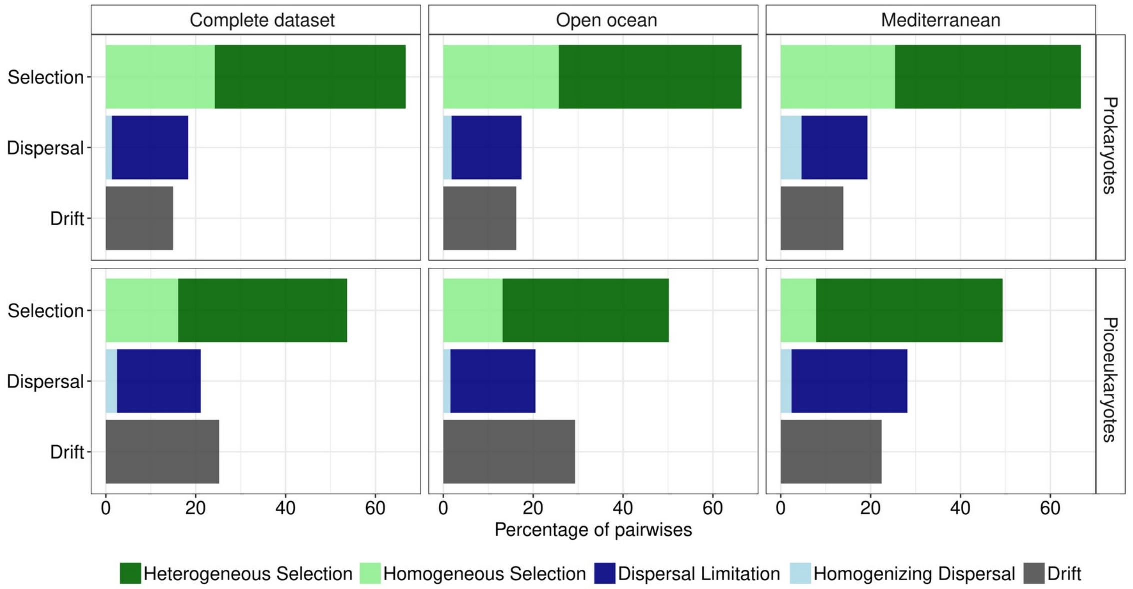


**Figure S8. Picoplankton community assembly processes integrating all depth zones.** Relative importance of the ecological processes structuring the picoplanktonic community using the complete dataset as well as separated by the open ocean and the Mediterranean Sea.

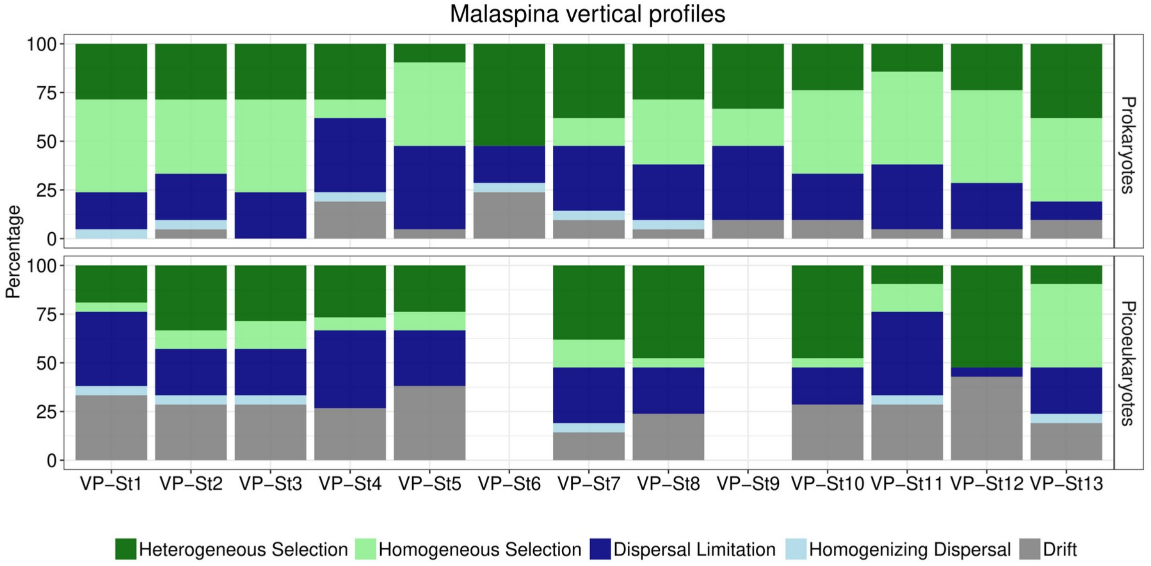


**Figure S9. Picoplankton community assembly processes in the *Malaspina* vertical profile stations.** Relative importance of the ecological processes structuring the picoplanktonic community integrating all depths (from 3 to 4000 m, i.e. 7 different depths) in each of the 13 vertical-profile (VP) stations (labeled as in Fig. 1A).

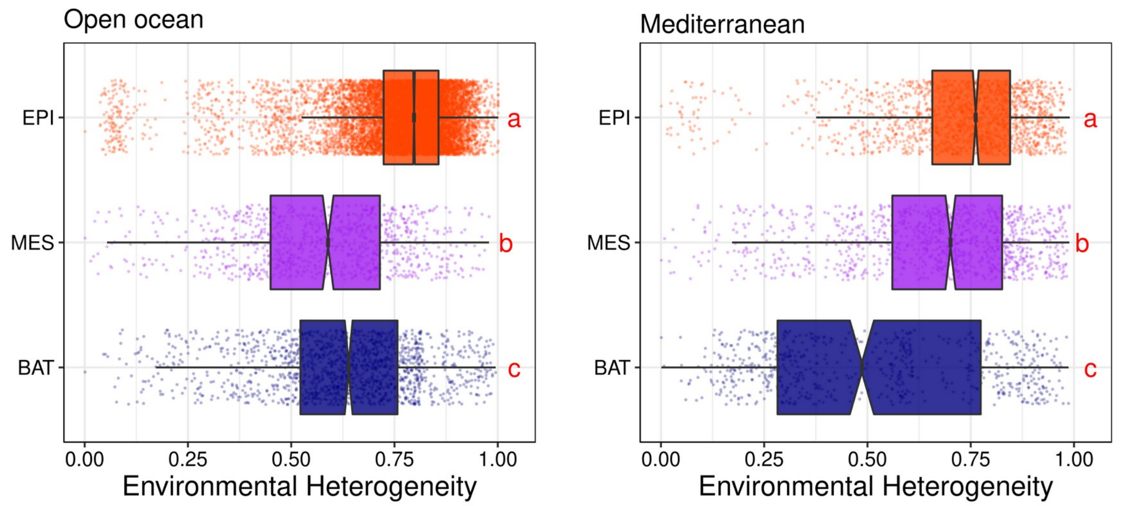


**Figure S10.** Environmental heterogeneity computed as the mean environmental dissimilarity between samples considering the main environmental variables (Temperature, Salinity, Fluorescence, $NO_3$, $PO_4$, $SiO_2$) in the open ocean and the Mediterranean Sea. Different red letters represent significantly different means [Kruskal-Wallis, Wilcoxon post-hoc test, $p<0.05$] between depth zones.

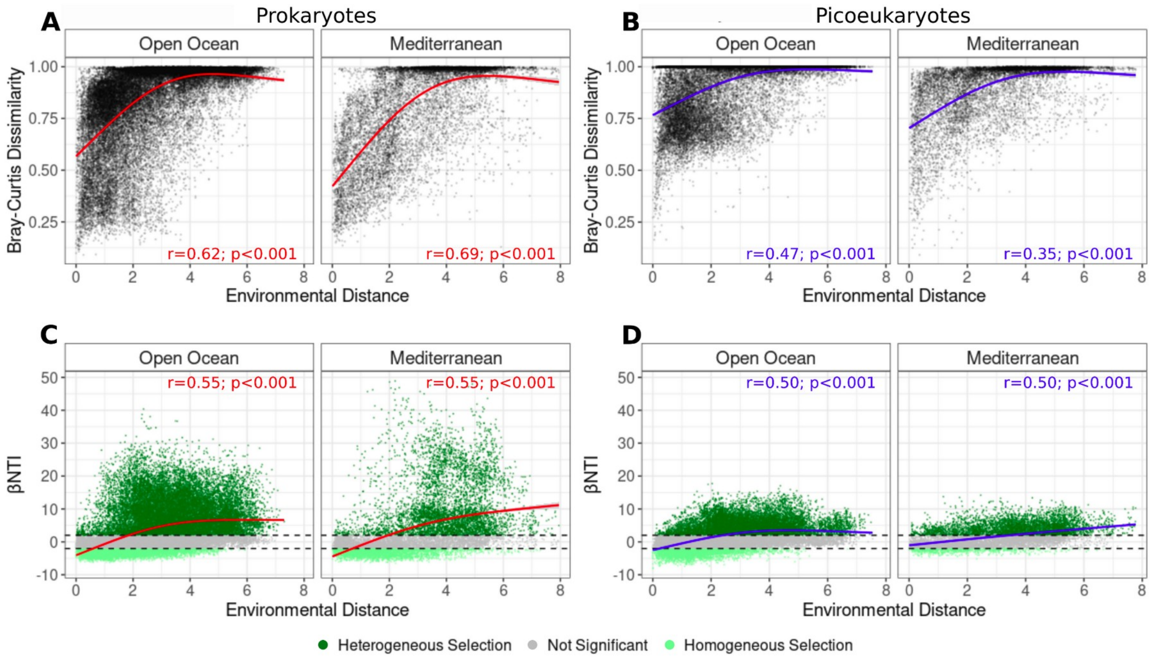


**Figure S11. Picoplankton community composition and phylogeny are positively related to environmental heterogeneity.** Difference in taxonomic (Bray-Curtis dissimilarity) and phylogenetic (βNTI) composition for all pairwise picoplankton community comparisons as a function of environmental distance for both prokaryotes **(A, C)** and picoeukaryotes **(B, D)** in the open ocean and Mediterranean Sea. The solid curves illustrate the nonlinear regressions. Spearman’s rank correlation coefficients are depicted on the panel. Outliers with high environmental distances (>10) corresponding to pairwise comparisons with epipelagic samples from the Costa Rica Dome upwelling system were removed from the open ocean plot (see Fig S13).

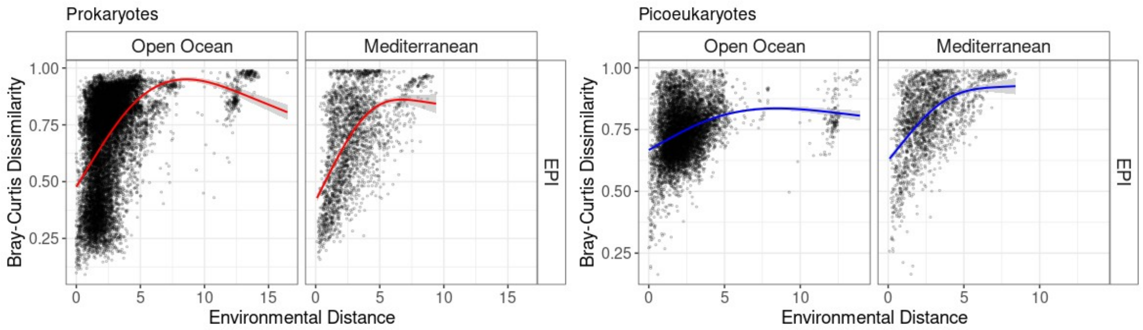


**Figure S12. Picoplankton community compositions are positively related to environmental heterogeneity.** Difference in composition (Bray-Curtis dissimilarity) for all pairwise picoplankton community comparisons as a function of environmental distance for both **(A)** prokaryotes and **(B)** picoeukaryotes in the epipelagic of the open ocean and Mediterranean Sea. The points with high environmental distances (>10) correspond to the pairwise comparisons with epipelagic samples from the Costa Rica Dome. The solid curves illustrate the nonlinear regressions. Spearman’s rank correlation coefficients are depicted on the panel.

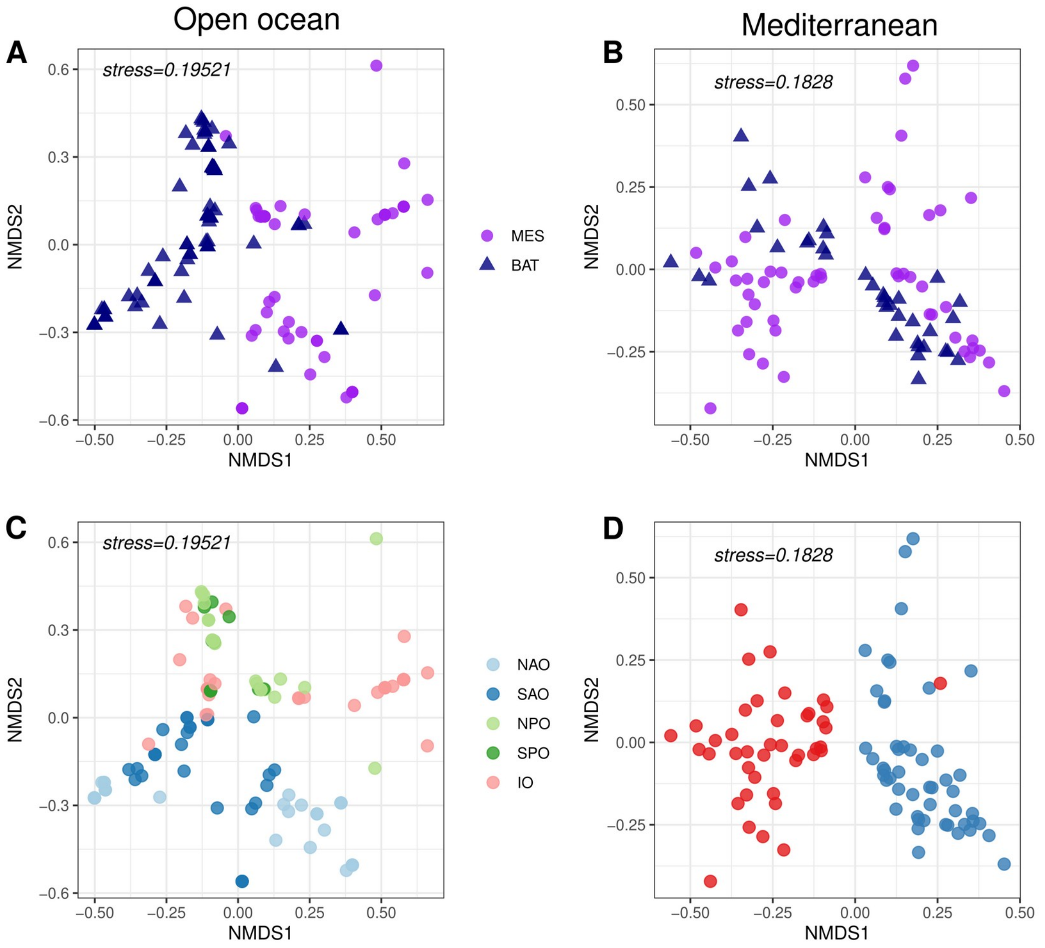


**Figure S13. Differences in water mass composition are segregated by depth zones and ocean basins.** Nonmetric multidimensional scaling (NMDS) based on the Euclidean distance of the samples' water mass composition – labeled by zones and basin – in the open ocean **(A, C)** and the Mediterranean Sea **(B, D)**. MES = Mesopelagic; BAT = Bathypelagic. NAO = North Atlantic Ocean, SAO = South Atlantic Ocean, NPO = North Pacific Ocean, SPO = South Pacific Ocean, IO = Indian Ocean, WMS = Western Mediterranean Sea, EMS = Eastern Mediterranean Sea.

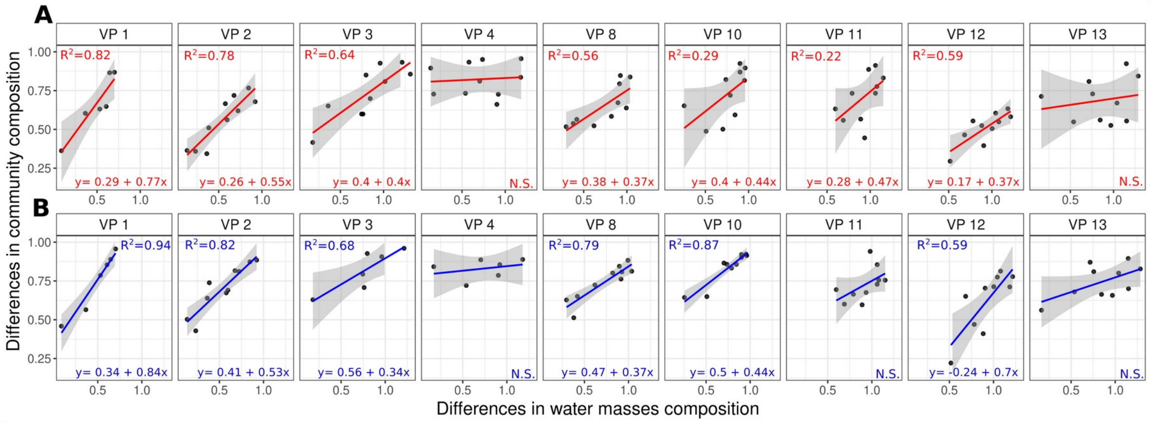


**Figure S14. Picoplankton community composition and potential dispersal are vertically linked to differences in water mass composition.** Difference in community composition (Bray-Curtis dissimilarity) as a function of water mass composition dissimilarity (Euclidean distances) for prokaryotes (in red) **(A)** and picoeukaryotes (in blue) **(B)** in *Malaspina* vertical profiles. Note that only meso- and bathypelagic samples were used in this analysis. The equation, the explanatory power of the linear regression models (adjusted R2), and the significance of the smooth terms (p<0.001) are shown on the plots.

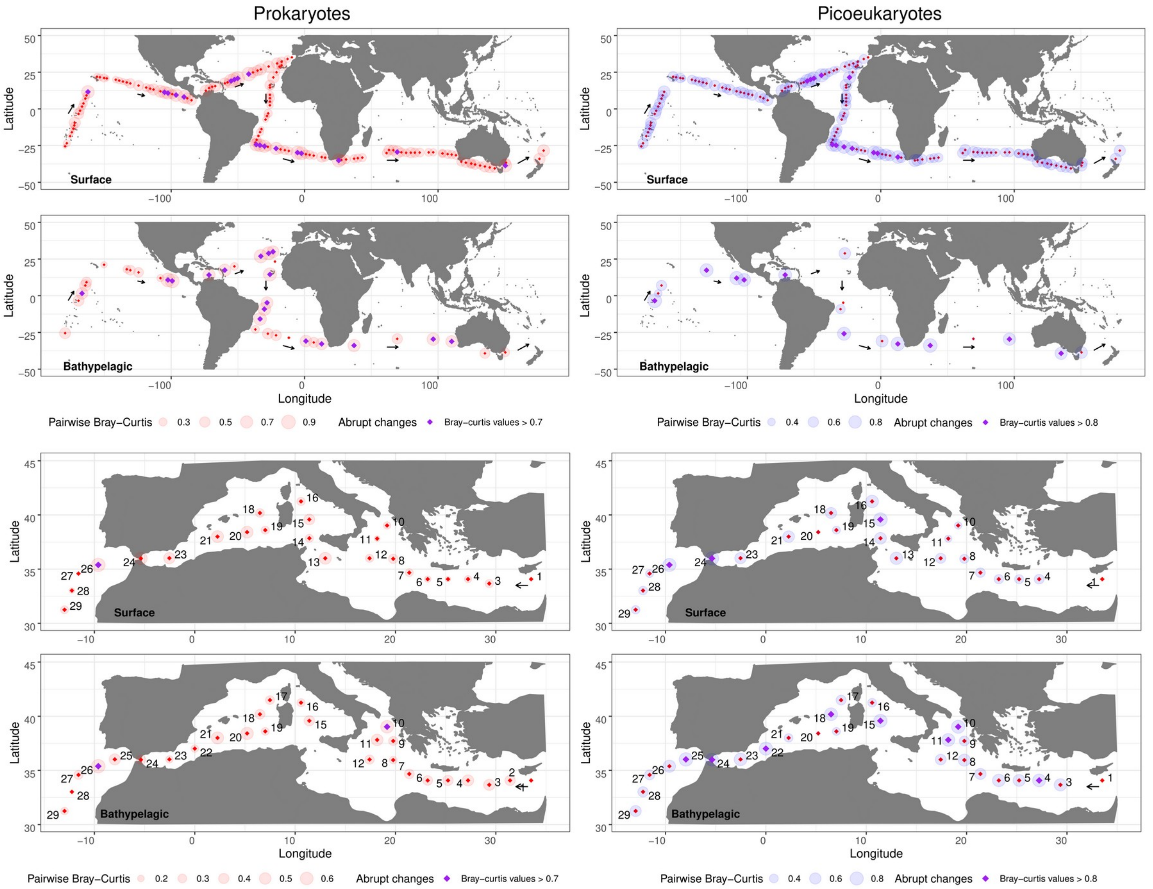


**Figure S15. Sequential change in community composition across space (sequential β-diversity).** Communities were sampled along the *Malaspina* and *Hotmix* expeditions (black arrows), and the composition of each community was compared against its immediate predecessor. The size of each bubble represents the Bray-Curtis dissimilarity between a given community and the community sampled previously.

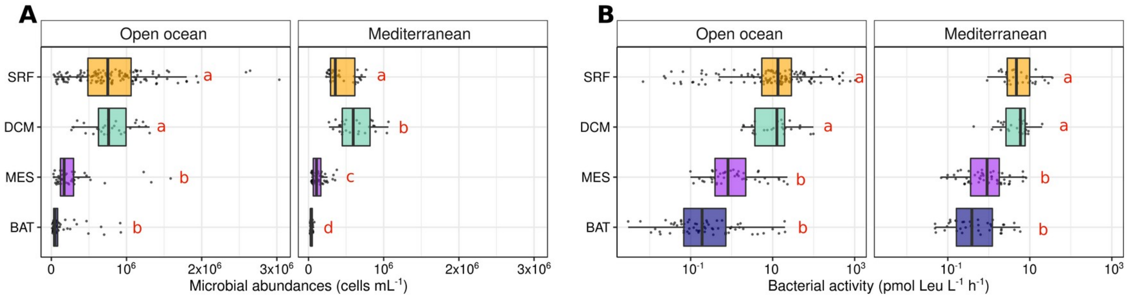


**Figure S16. Microbial abundances and bacterial activity sharply decrease in deep waters**. **(A)** Microbial abundances (prokaryotes + picoeukaryotes) as measured by flow cytometry; **(B)** bacterial activity as measured by leucine incorporation rates in each zone (SRF, surface; DCM, deep chlorophyll maxima; MES, Mesopelagic; BAT, Bathypelagic) of the open ocean and the Mediterranean Sea. Different red letters represent significantly different means [ANOVA, Tukey post-hoc test, $p<0.05$] between depth zones.
